# Alternative cell entry mechanisms for SARS-CoV-2 and multiple animal viruses

**DOI:** 10.1101/2023.07.02.547368

**Authors:** Ravi Ojha, Anmin Jiang, Elina Mäntylä, Naphak Modhira, Robert Witte, Arnaud Gaudin, Lisa De Zanetti, Rachel Gormal, Maija Vihinen-Ranta, Jason Mercer, Maarit Suomalainen, Urs F. Greber, Yohei Yamauchi, Pierre Yves-Lozach, Ari Helenius, Olli Vapalahti, Paul Young, Daniel Watterson, Frédéric A. Meunier, Merja Joensuu, Giuseppe Balistreri

## Abstract

The cell entry mechanism of SARS-CoV-2, the causative agent of the COVID-19 pandemic, is not fully understood. Most animal viruses hijack cellular endocytic pathways as an entry route into the cell. Here, we show that in cells that do not express serine proteases such as TMPRSS2, genetic depletion of all dynamin isoforms blocked the uptake and strongly reduced infection with SARS-CoV-2 and its variant Delta. However, increasing the viral loads partially and dose-dependently restored infection via a thus far uncharacterized entry mechanism. Ultrastructural analysis by electron microscopy showed that this dynamin-independent endocytic processes appeared as 150-200 nm non-coated invaginations and was efficiently used by numerous mammalian viruses, including alphaviruses, influenza, vesicular stomatitis, bunya, adeno, vaccinia, and rhinovirus. Both the dynamin-dependent and dynamin-independent infection of SARS-CoV-2 required a functional actin cytoskeleton. In contrast, the alphavirus Semliki Forest virus, which is smaller in diameter, required actin only for the dynamin-independent entry. The presence of TMPRSS2 protease rescued SARS-CoV-2 infection in the absence of dynamins. Collectively, these results indicate that some viruses such as canine parvovirus and SARS-CoV-2 mainly rely on dynamin for endocytosis-dependent infection, while other viruses can efficiently bypass this requirement harnessing an alternative infection entry route dependent on actin.

## Introduction

To gain access into the cytoplasm of their host cells, where synthesis of viral proteins and genome replication occur, most animal viruses hijack the endocytoic pathway^1^ by which the cell injests particles and nutrients by encoulfing them into vesicular compartments. Multiple endocytic mechanisms have been described and viruses have been shown to utilize one or more of these pathways^3^. In addition to endocytosis, some viruses have been show to infect cells by penetrating directly through the plasma membrane (PM)^2^. Understanding how viruses utilise these processes is key to the identification of potential cellular targets that can be exploited to block viral infections, and to increase our knowledge of key cellular processes involved in the viral entry and infection that are currently incompletely understood.

The final step of an endocytic vesicle formation culminates with the pinching of vesicle off from the PM into the cytoplasm. This fission process requires energy which is generated by one of the three dynamin (dyn1-3) isoforms, that all possess GTPase activity^4^. Not all endocytic pathways require dynamin and, to complete the PM invagination process and generate endocytic vesicles, cells can use the force generated by the cooperation of the actin cytoskeleton, membrane deforming proteins, and motor proteins^5–7^. Virions taken up by endocytosis, are sorted to intracellular, membrane enclosed organelles from which the viral genome ‘escapes’ and reaches the protein synthesis and replication machineries present in the cytoplasm and/or nucleus of the host cell to replicate.

For enveloped viruses, i.e., viruses surrounded by lipid membranes, the delivery of viral genome into the cytoplasm is mediated by the fusion of the viral and cellular membranes, a process driven by viral surface proteins (often referred to as spikes on the virion). For some enveloped viruses the cue that triggers fusion is the drop in pH that occurs once the viral particle reaches the lumen of endosomal vesicles (e.g., Influenza virus, Vesicular stomatitis virus, Semliki Forest virus, among others)^8^. For other viruses, such as respiratory viruses (including coronaviruses), the fusion is triggered by proteolytic cleavage of the spike proteins that, once cleaved, undergo conformational changes leading first to the insertion of the viral spike into the host membrane and, upon retraction, the fusion of viral and cellular membranes^9,10^. In the case of coronaviruses, these proteolytic cleavages are performed by cellular proteases present either in the endo-lysosomal compartment (e.g., the cysteine protease cathepsin-L) or at the cell surface (e.g., the transmembrane serine protease 2, TMPRSS2)^11–14^. Depending on the availability of these proteases, virus fusion and the delivery of the viral genome into the cytoplasm can either occur at the PM (i.e., in cells that express PM serine proteases) or from within endo-lysosomes (i.e., in cells that express active cathepsins in endo/lysosomes but not serine proteases at the cell surface)^15^. In the latter case, the virus requires endocytosis to reach endosomes and lysosomes. This route of cell entry has been proposed for the current SARS-CoV-2 variant of concern (VOC) Omicron, the infection of which seems to favour the endo/lysosomal entry route rather than direct fusion at the PM^16^. In addition, SARS-CoV-2 infection of human neurons also seems to require endocytosis and endosomal transport^17^.

Here, we surveyed a range of animal viruses for their dynamin-dependency to infect cells. We show that while some viruses, including SARS-CoV-2, strongly depend on the presence of dynamins to productively infect cells, other animal viruses, including alphaviruses, influenza, and bunyaviruses, among others, can use dynamin-independent endocytosis (DIE) as an alternative and efficient entry mechanism. This yet uncharacterized endocytic pathway is sensitive to actin perturbations and is morphologically distinct from other well characterized endocytic processes.

## Results and discussion

### Characterization of dynamin 1,2 conditional knock-out cells to study virus entry

Many animal viruses, including the alphaviruses Semliki Forest (SFV) and Sindbis (SINV) virus, influenza virus (IAV), and vesicular stomatitis virus (VSV), have been shown to infect cells using dynamin-dependent endocytosis^3^. Most of these studies have been performed by treating cells with small molecule dynamin inhibitors, by overexpression of dominant-negative inactivated forms of dynamin, or by depletion of dynamin mRNA levels using RNA interference methods. These loss-of-function approaches, although easy to implement, suffer from uncharacterized off-targets effects^18^ or incomplete depletion of dynamin isoforms. To address the role of dynamins in virus infection, and overcome the above-mentioned limitations, we started our investigation by using genetically engineered dynamin 1,2 conditional double knockout (KO) mouse embryonic fibroblasts (MEF^DNM1,2 DKO^)^19^. In these cells, the two main isoforms of dynamin (dyn1,2) can be completely depleted within 6 days by the addition of 4OH-tamoxifen (4OH-TMX)^19^ (Figure 1 A). The third isoform of dynamin (dyn3) is not detectable in these cells^19^. To functionally monitor the specificity and level of inhibition of dynamin-dependent endocytosis in this model system, we used transferrin (Tf) as positive control, which is internalized in a dynamin-dependent manner^20^, and the B subunit of the cholera toxin (CTB), which can enter cells via both dynamin-dependent^21,22^ (at low doses) or dynamin-independent (at high doses) endocytic mechanisms^23^. As expected, 6 days after 4OH-TMX treatment, the uptake of fluorescently labelled Tf (5 μg/ml) was dramatically inhibited in most cells (Figure 1 B). Consistent with previous a report ^19^, in all experiments approximately 3-5% of the MEF^DNM1,2 DKO^ cells did not respond to 4OH-TMX treatments and as a result maintained normal levels of Tf uptake (Figure S1). The internalization of CTB was also significantly reduced at low toxin concentration (i.e. 50 ng/ml) (Figure 1 C), while at higher concentration (i.e. 1 μg/ml), the levels of CTB uptake in dyn1,2 depleted cells were comparable to those of MEF^DNM1,2 DKO^ cells that were treated with vehicle control (Figure 1 D). These experiments confirmed that the MEF^DNM1,2 DKO^ represent a suitable model system to study dynamin-dependent and -independent endocytosis.

**Figure 1.**
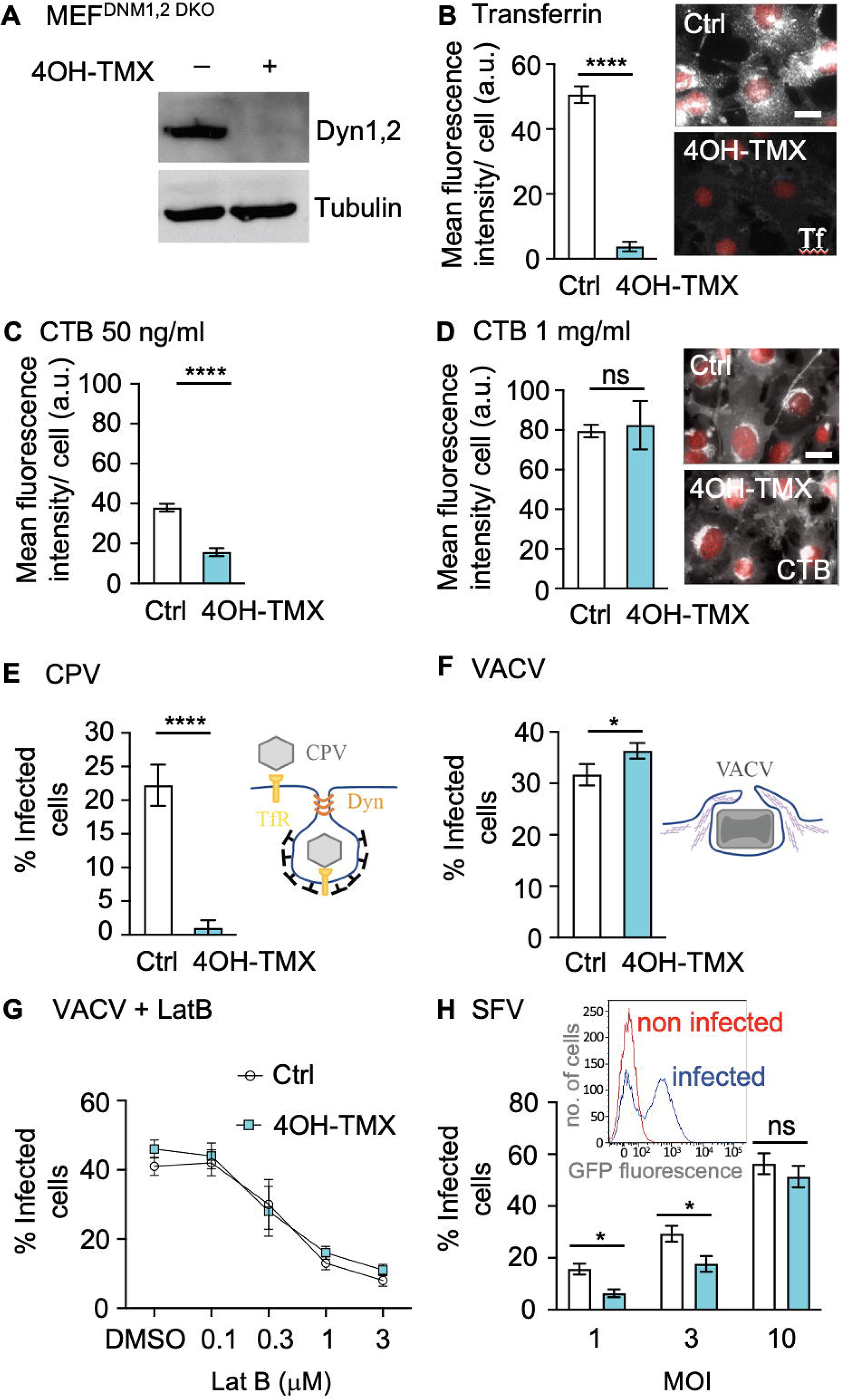
Characterization of MEF dynamin 1,2 conditional KO cells to study virus infection. A) Western blot analysis of dynamin 1 and 2 levels in MEF^DNM1,2, DKO^ cells treated with vehicle control or 4OH-TMX for 6 days. Tubulin was used as a loading control. The Dyn1,2 antibody used recognizes both dynamin 1 and 2. B) Quantification of Tf Alexa-647 (Tf) (1 μg/ml) 30 min uptake in MEF ^DNM1,2, DKO^ cells treated with vehicle control or 4OH-TMX for 6 days prior fixation and Hoechst DNA staining. Representative fluorescent images on the right show Tf (white) and Hoechst (red) in MEF^DNM1,2, DKO^ cells treated with vehicle control or 4OH-TMX. C-D) Quantification of CTB Alexa-647 (CTB) uptake in MEF^DNM1,2, DKO^ cells treated with vehicle control or 4OH-TMX for 6 days and incubated with indicated concentrations of CTB for 30 min prior to fixation and Hoechst DNA staining. Representative fluorescent images on right show CTB (white) and Hoechst (red) in MEF^DNM1,2, DKO^ cells treated with vehicle control or 4OH-TMX. E) Quantification of CPV infection in MEF^DNM1,2, DKO^ cells treated with vehicle control or 4OH-TMX for 6 days and infected with CPV for 24h, and immunostained for non-structural protein 1 (NS1). The inset schematic illustrates the entry mechanism of CPV. F) Quantification of VACV infection in MEF^DNM1,2, DKO^ cells treated with vehicle control or 4OH-TMX for 6 days and infected with VACS for 6 h. The inset schematic illustrates VACV entry by micropinocytosis. G) Effect of Latrunculin-B (LatB) on VACV infection in MEF^DNM1,2, DKO^ cells treated with vehicle control or 4OH-TMX for 6 days and infected with VACS for 6 h. H) Quantification of SFV infection in MEF^DNM1,2, DKO^ cells treated with vehicle control or 4OH-TMX for 6 days and infected with indicated MOIs of SFV-EGFP for 7 h. The inset illustrates the FACS analysis of virus-induced EGFP fluorescence in non-infected (red line) and infected (blue line) vehicle-control treated cells. Values represent the mean of 3 independent experiments. Error bars represent the standard deviation (STDEV). Statistical significance was calculated using a unpaired two-tailed t-test (*p<0,05; ****p<0,0001; n.s.= non-significant).

To test whether the inducible dynamin depletion system was suitable to study virus infection, we first tested three well characterized viruses in the MEF^DNM1,2 DKO^ cells treated with vehicle-control or 4OH-TMX for 6 days: i) canine parvovirus (CPV), a small single-stranded DNA virus that uses the Tf receptor (TfR) and dynamin-dependent endocytosis to enter cells^24,25^ (here used as a positive control); ii) vaccinia virus (VACV-EGFP, EGFP expressed under viral early gene promoter^26^), a large DNA virus that enters cells via actin-dependent, dynamin-independent macropinocytosis^27,28^ (negative control); and Semliki Forest virus (SFV-GFP, GFP expressed as a fusion with viral replicase protein nsP3^29^), which has been shown to infect cells mainly via dynamin-dependent endocytosis^8^, although alternative entry mechanisms have also been proposed^30^. Infection rates were determined by Fluorescence-activated cell sorting (FACS) flow cytometry analysis of virus-induced expression of GFP or after immunofluorescence staining of viral proteins. Depletion of dyn1,2 in MEF^DNM1,2 DKO^ cells transiently over-expressing the feline TfR^25^ inhibited CPV infection by more than 90% (Figure 1 E and S2). The infectivity of VACV in dyn1,2 depleted MEF^DNM1,2 DKO^ cells was comparable to that of control cells, confirming that this virus enters cells via a dynamin-independent mechanism (Figure 1 F). Disruption of the actin cytoskeleton using the actin depolymerizing drug Latrunculin-B (LatB), on the other hand, blocked VACV infection in both dyn1,2 depleted and control MEF^DNM1,2 DKO^ cells (Figure 1 G). The actin-dependency of VAVC infection was expected and consistent with entry by macropinocytosis, a process where actin polymerization is required to support the formation of PM protrusions that can engulf extracellular fluids and large particles such as VACV particles that have a size of approximately 250×250×350 nm^27,28,31^. Notably, the infection of SFV was only partially inhibited by dyn1 and 2 depletion, and increasing the virus dose (i.e., the multiplicity of infection, MOI=10), fully restored infection (Figure 1 H).

In summary, the MEF^DNM1,2 DKO^ cells are a suitable model to study the role of dynamin in virus infection. Viruses that use a receptor that is internalized by dynamin-dependent endocytosis (e.g. CPV and the TfR) cannot efficiently infect cells in the absence of dynamins. Compensatory dynamin-independent endocytosis does not appear to be available for these ‘single-receptor’ viruses. On the other hand, viruses such as SFV, which may use more than one protein receptor, e.g. a variety of heparan-sulphate-containing glycoproteins^32^, are internalized primarily by dynamin-dependent endocytosis, but if this entry mechanism is not available, infection can still occur via an alternative dynamin-independent pathway.

### Dynamin-independent endocytosis is an alternative, efficient entry pathway for multiple animal viruses

Endocytic pathways that require dynamin, such as clathrin-mediated endocytosis (CME), have been associated with the cell entry of numerous viruses, including members of the mosquito-delivered alphaviruses, such as SFV and SINV^33^, IAV^34^, as well as members of the *Bunyaviridae* such as Uukuniemi virus (UUKV)^35^, the vesiculoviruses such as vesicular stomatitis virus (VSV)^36,37^, common cold human rhinovirus (RV) B14 and A89^33,38^, and species C AdV such as AdV-C2 or C5^40–43^ but also AdV-B3 and AdV-B35, although the dynamin requirement of B3 and B35 was variable between cell lines ^44,45^. The results obtained with SFV (Figure 1 H) prompted us to test if the dynamin-independent pathway could be used as a productive entry route for other viruses that are known to use dynamin-dependent endocytosis to enter host cells. To this end, we tested if the infectivity of SINV, VSV, IAV, UUKV, HRV-A1 and AdV-C5 was inhibited in dyn1,2 depleted MEF^DNM1,2 DKO^ cells. VACV, which is known to infect cells by actin-dependent macropinocytosis irrespective of the presence of dynamins, was used as a comparison (Figure 1 F-G). Infection was monitored by FACS or immunofluorescence analysis. VACV^26^, VSV^37^, and AdV-C5^39^ were engineered to express the EGFP protein, and SINV to express the mCherry, as fluorescent reporters of infection. MEF^DNM1,2, DKO^ cells were treated with vehicle control or 4OH-TMX for 6 days, and the infection rates for these recombinant viruses were quantified at 7 hours post infection (hpi) (VACV-EGFP, VSV-EGFP, SINV-mCherry) and 22 hpi (AdV-C5-EGFP) by monitoring the EGFP expression, or by using immunofluorescence staining against viral antigens with virus-specific antibodies at 8 hpi (IAV X31, UUKV) and 22 hpi (HRV-A1). Interestingly, all the tested viruses, albeit with different efficiency, were able to infect cells in the absence of dynamins (Figure 2). At low viral doses (equivalent to an infection of approximately 20-40% of cells), the infections of SINV-mCherry, VSV-EGFP, and IAV X31 were significantly decreased in dyn1,2 depleted cells. In contrast, in the absence of dynamins, the infection of UUKV was not significantly blocked, and in the case of the common cold viruses HRV-A1 and AdV-C5-EGFP, infection in dynamin-depleted cells was enhanced in comparison to controls (Figure 2 and S3). Thus, dynamin-independent endocytosis is an efficient, alternative virus entry portal that can be exploited by multiple animal viruses. The results also demonstrate that using a low dose of virus is advisable to estimate the contribution to infection of the two entry mechanisms. A comparative analysis of all tested viruses, at a viral dose that corresponds to 20-40% infection rates in vehicle control treated cells (Figure 2 and S2), indicates that the dependence on dynamin-mediated endocytosis follows this qualitative order: VSV>SFV=SINV=IAV>UUNV. Infections with VACV, RV-A1a and AdV-C5 were not inhibited by dynamin depletion, rather increased, suggesting that the Dyn-KO cells may have adapted by up-regulation of dynamin-independent endocytic pathways. Additionally, the permanent knockout of dynamins may impact on other processes than membrane fission at the plasma membrane, e.g., the contractile machinery at the interface between cytoplasmic membranes and the actin cytoskeleton ^47^, or the suppression of tubulin acetylation promoting dynamic instability of microtubules ^48 49^.

**Figure 2.**
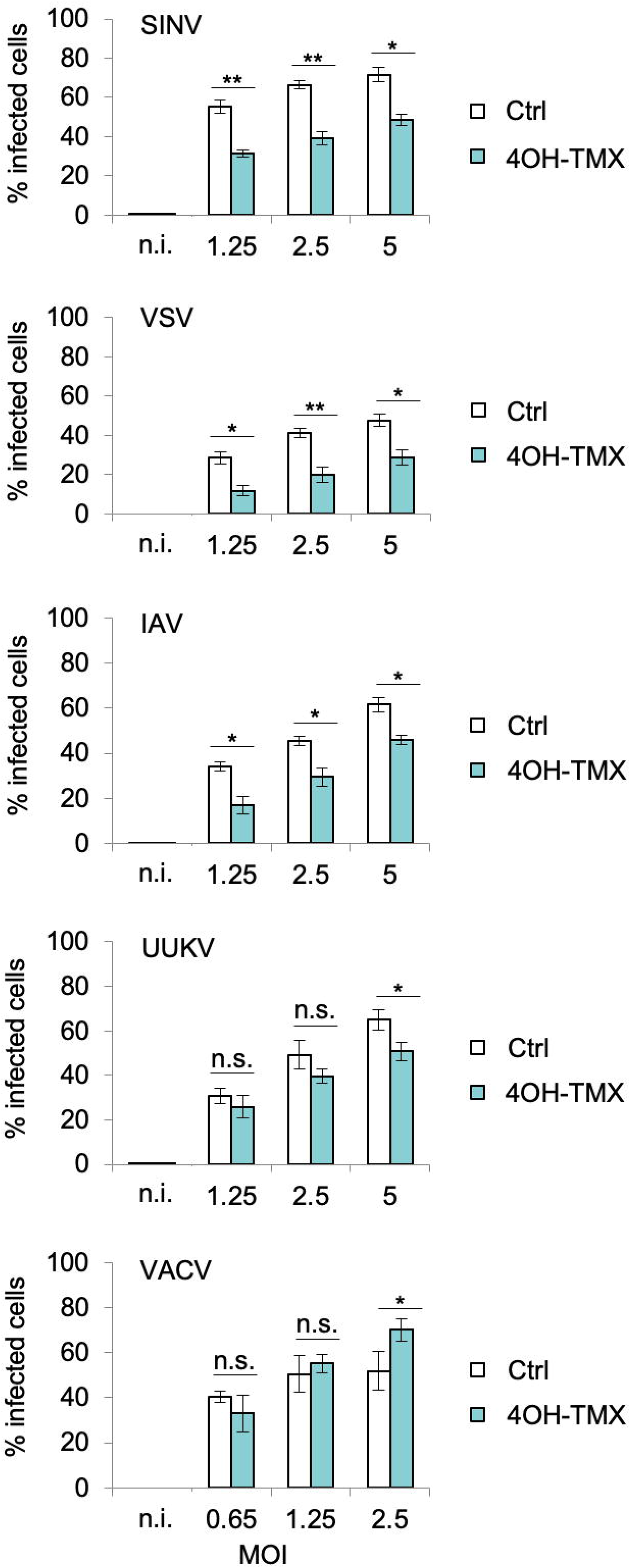
Dynamin-independent endocytosis is an alternative, efficient virus entry pathway for multiple animal viruses. Infection of indicated viruses in MEF^DNM1,2, DKO^ cells treated with vehicle control or 4OH-TMX for 6 days and infected for 6 h (VACV), 7 h (SINV, VACV), and 8 h (IAV X31, UUKV). Virus infection was determined by FACS analysis of virial induced EGFP (VAVC and VSV), mCherry (SINV) or after immunofluorescence of viral antigens using virus-specific antibodies (IAV X31 and UUKV). Values indicate the mean of three independent experiments and the error bars represent the standard deviation (STDEV). n.i.= non infected. Statistical significance was calculated by unpaired two-tailed t-test (*p<0,05; ** p<0,01; n.s.= non-significant).

### Dynamin-independent virus entry is facilitated by the actin cytoskeleton

Dynamin-depleted cells were efficiently infected by SFV at the higher virus doses (Figure 1 H). A set of small molecule inhibitors, known to block different endocytic pathways, was used to further characterized the dynamin-independent endocytic mechanism used by SFV. The drugs were added to the MEF^DNM1,2 DKO^ cells (pre-treated for 6 days with vehicle control or 4OH-TMX) 15 min before infection and were present throughout infection for 1 h. To prevent further SFV cytosolic entry following removal of virus inoculum, cells were maintained in media containing NH_4_Cl, which neutralized endosomal pH (Figure 3 A). Both in control and dyn1,2 depleted cells, SFV infection was highly sensitive to hyper-osmotic shock induced by 0.45 M sucrose, a treatment that blocks endocytosis, and drugs known to inhibit endosomal acidification, i.e. Bafylomycin A1 (BafA) and NH_4_Cl (Figure 3 B). Thus, even in the absence of dynamin, SFV infection required endocytosis and, as described earlier, delivery of viral particles to acidic intracellular endosomes^8^. Unexpectedly, inhibitors of CME (e.g. chlorpromazine)^40^, and dynamins^42^ (i.e. dyngo-4A and Dynole^43^) blocked infection to a similar extent in both control as well as dyn1,2 depleted cells, indicating potential off-target effects of these drugs. Incubation of cells with the phospho-inhositol 3 kinase (PI3K) inhibitor wortmannin, Rac1/CdC42 GTPase inhibitor ML141, or at lower temperatures (i.e. 24 °C), three treatments that are known to specifically block macropinocytosis, were either ineffective (wortmannin) or moderately effective (ML142 and low temperature) to a similar extent in both control and dyn1,2-depleted cells, indicating that these endocytic processes are different from *bona fide* macropinocytosis.

**Figure 3.**
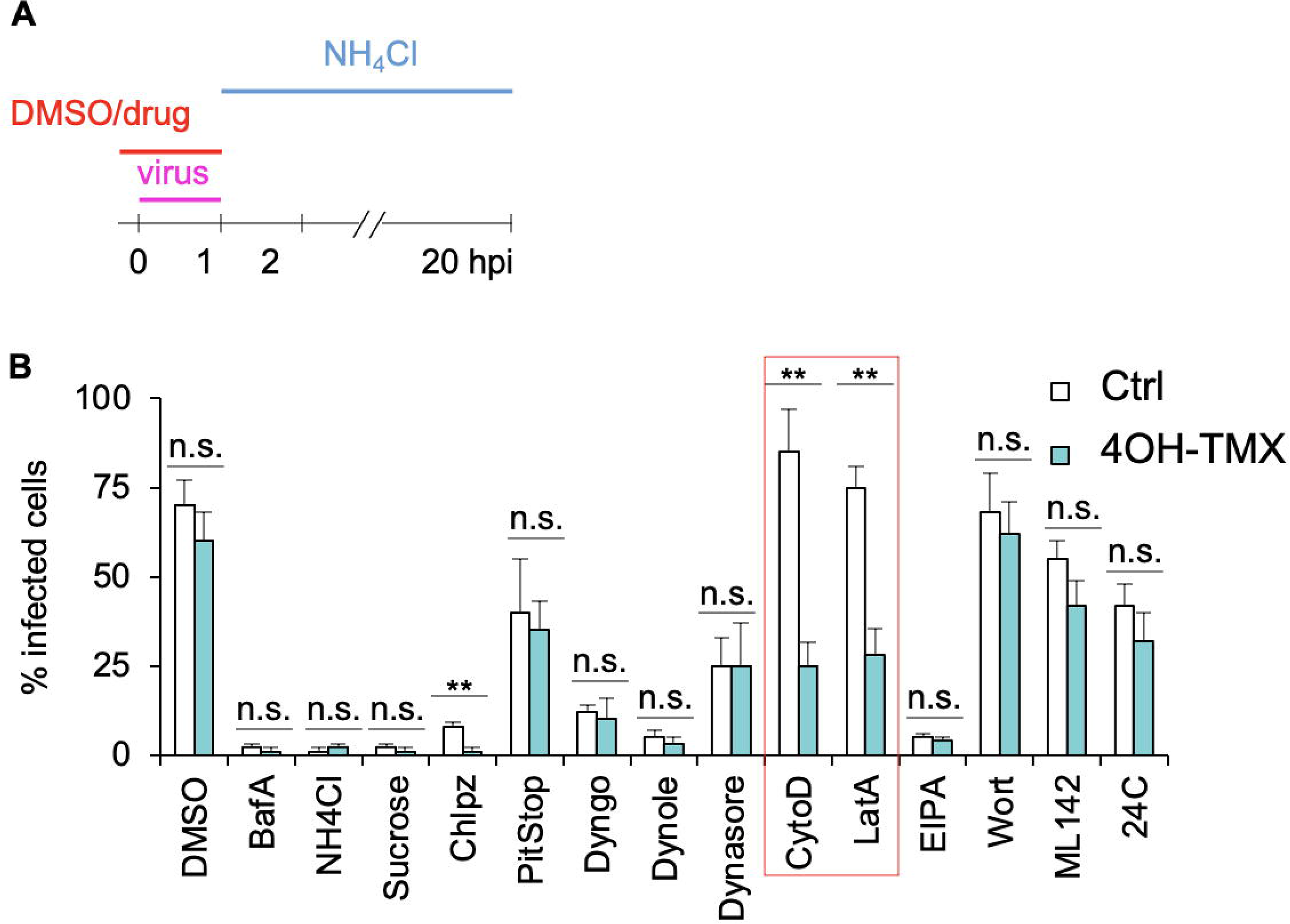
Dynamin-independent virus entry is actin dependent. A) Schematic description of indicated drug treatments in MEF^DNM1,2 DKO^ cells following 6-day treatment with vehicle (ctrl) or 4OH-TMX, and infected with SFV-EGFP. B) Quantification of the experiment described in A. After drug treatment and infection, the percentage of infected cells was determined by FACS analysis of virus induced expression of EGFP. The red-boxed area indicates the treatments with actin depolymerizing drugs. Values indicate the mean of three independent experiments and the error bars represent the standard deviation. Statistical significance was calculated by unpaired two-tailed t-test (** p<0,01; n.s.= non-significant).

Treatments with actin depolymerizing agents such as cytochalasin-B (CytoD) and Lat-A, did not inhibit SFV infection in control MEF^DNM1,2 DKO^ cells (Figure 3 B, Ctrl bars). Thus, unlike the large VACV, the smaller SFV (approximately 60 nm in diameter)^44^ does not require functional actin cytoskeleton for the entry process in unperturbed cells. However, in dyn1,2-deficient MEF^DNM1,2 DKO^ cells, the treatment with CytoD and Lat-A and significantly reduced SFV infection (Figure 3 B) by approximately 60% as compared to control cells treated with the same drugs (Figure 3 B, red boxed area). Hence, dynamin-dependent and -independent endocytic pathways differ in their sensitivity to actin-depolymerizing drugs.

To further study the differences in the actin cytoskeleton in control and dyn1,2-depleted cells, we performed single-molecule resolution total internal reflection fluorescence microscopy (TIRFM) in live MEF^DNM1,2 DKO^ transiently expressing LifeAct-mEos2, a recombinant fusion protein of LifeAct^45^ (labels actin fibres) and the photocovertible mEos2^46^ (Figure S4 A-B). Firstly, to test if this imaging system could be used to quantitatively monitor subtle, global changes of cellular actin dynamics, we treated MEF^DNM1,2 DKO^ cells with low doses of Lat-A (0.1 μM) or vehicle control for 10 min. Both the frequency distribution of diffusion coefficients (Log_10_D, where [D] = µm^2^ s^−1^) and the mean square displacement (MSD; µm^2^) of single-molecules of LifeAct-mEos2 decreased in Lat-A treated cells, indicating an overall slower mobility of LifeAct-mEos2-labelled actin fibers (Figure S4 C). The effect of dyn1,2 depletion on the overall mobility of LifeAct-mEos2 labelled actin was comparable to that of low doses of Lat-A, indicating similar change in actin dynamics (Figure S4 D). Both the dyn1,2 depletion with 4OH-TMX and the treatment with Lat-A induced similar changes in the actin cytoskeleton of the MEF cells, resulting in an accumulation of short and puncatate filament structures that were observed with phalloidin staining (Figure S4 F).

### Ultrastructural analysis of dynamin-independent virus entry

Because SFV entered cells efficiently both in the presence and absence of dynamins, we used transmission electron microscopy (TEM) to gain ultrastructural information on endocytic processes that mediate SFV entry in dyn1,2 depleted cells. MEF^DNM1,2 DKO^ cells were treated with 4OH-TMX or vehicle control for 6 days and, following virus (MOI=1000) adsorption at 4°C for 1 h, cells were shifted at 37 °C for 10 minutes to promote viral internalization. In MEF^DNM1,2 DKO^ cells treated with vehicle control, TEM analysis revealed numerous viruses at the outer surface of the cells (Figure 4 A), as well as inside endocytic invaginations that appeared surrounded by an electron dense coated, consistent with the appearance of clathrin coated pits^47,48^ (CCP) (Figure 4 B). A small, but sizable fraction of SFV particles were also found inside bulb-shaped non-coated pits (NCP) (Figure 4 C). Occasionally, large (>1µm in diameter) irregular vacuoles located close to the PM and containing viruses were detected. These structures, here annotated as ‘large endocytic profiles, LEP’, could represent either macropinocytic processes or slight invaginations of the PM that appear circular after TEM cross-sectioning (Figure 4 D). In dyn1,2 depleted MEF^DNM1,2 DKO^ cells, SFV virions were also readily detected at the PM (Figure 4 F). CCPs were often associated with long tubular membranous structures (Figure 4 G). In agreement with these results, it had been reported that in the absence of dynamin the final step of CME does not occur, and the respective vesicles are pulled from the PM towards the cytoplasm by actin polymerisation followed by depolymerization and release back towards the PM^19^. A fraction of the viruses was found trapped in these ‘stalled’ CCPs (Figure 4 G), and the rest in NCPs (Figure 4 H) and in LEPs (Figure 4 I). Quantitative analysis of the TEM data revealed that after dyn1,2-depletion, the most abundant virus-containing endocytic processes were the NCPs (Figure 4 I), that reached a relative abundance of 41.7% of all virus-containing invaginations.

**Figure 4.**
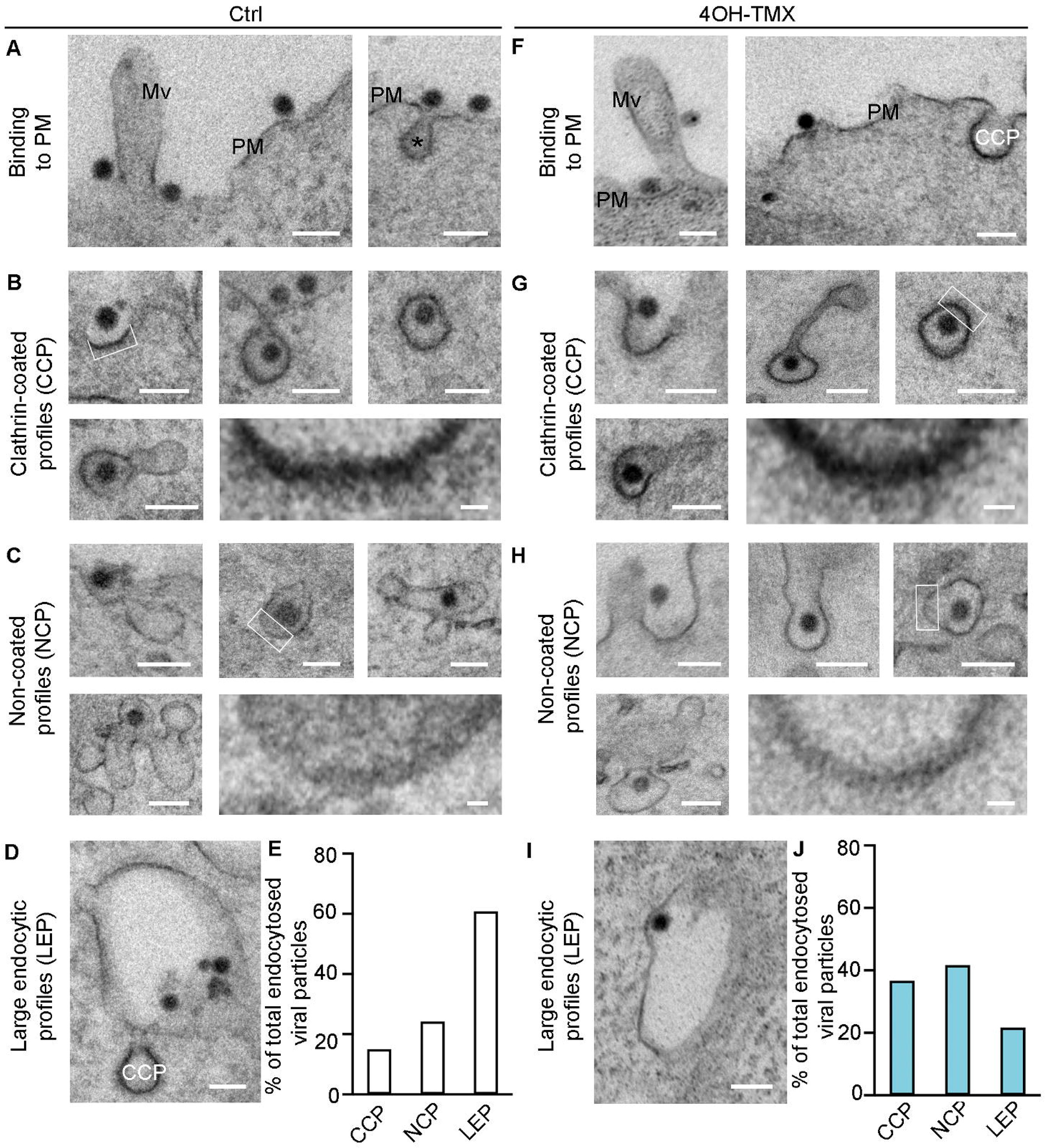
Ultrastructural analysis of dynamin-independent virus entry. Representative TEM images of MEF^DNM1,2, DKO^ cells treated with vehicle control (A-D) or 4OH-TMX (E-I) for 6 days and infected with SFV (MOI 1000) on ice for 1.5 h followed by shift to 37°C for 10 min before fixation and processing for TEM. The fraction of total viral particles found in each of the described endocytic processes is quantified in E and J, respectively. Scale Bar 100 nm. CCP = clathrin-coated pit, Mv = microvilli, PM = plasma membrane, asterisks (*) indicate stalled endocytic pits. Boxed areas are magnified in bottom right corner of each figure panel. Quantification of each treatment (EtOH vehicle ctrl or 4OH-TMX) includes over 500 viral particles from 2 independent experiments.

This analysis is consistent with two entry mechanisms for SFV, one clathrin- and dynamin-dependent, and another clathrin- and dynamin-independent. We speculate that upon depletion of dynamins, a fraction of the virus remains trapped in CCP. This stalled entry pathway might account for the partial inhibition of infection in Dyn-KO cells observed for SFV and possibly for other viruses. The fact that in unperturbed cells, SFV infection is not sensitive to actin depolymerizing drugs, suggests that the dynamin-dependent route is the predominant entry pathway. However, a sizable fraction of the virions can access an alternative entry pathway that, based on the effect of actin depolymerizing drugs, seems to rely more on the actin cytoskeleton. The precise role of actin is unknown but it could potentially help the final pinching step of the dyn-independent endocytic process.

### ACE2-mediated endocytosis of SARS-CoV-2 trimeric spike proteins is dynamin-dependent

The exact mechanism of SARS-CoV-2 cell entry is not fully understood, and evidence suggests that both clathrin-dependent and -independent endocytosis could be involved in cell culture models. A study from Bayati *et al*.^49^ used chemical inhibitors of dynamins and siRNA-mediated depletion of clathrin to show that the internalization of both the viral trimeric soluble Spike protein of SARS-CoV-2 and the infection of lentiviruses pseudotyped with SARS-CoV-2 S were decreased. Another study indicated that cell entry of SARS-CoV-2 spike protein occurs via clathrin independent endocytosis in cells devoid of the human angiotensin converting enzyme 2 (ACE2)^50^. Both studies implied entry via dynamin-dependent mechanisms.

To address the role of endocytosis in SARS-CoV-2 infection, and to identify the potential endocytic mechanisms that leads to productive entry, we started our investigations by following the uptake and intracellular trafficking of fluorescently labelled, recombinant soluble trimeric spike proteins (S) of SARS-CoV-2. This soluble version of the viral spike protein is held as a trimer by a molecular clamp^51,52^ that replaces the trans-membrane domain of the glycoprotein. In addition, for stabilization purposes, the polybasic furin-cleavage site, required to render the viral spike fusogenic, was rendered uncleavable in each monomer by mutagenesis^49^. MEF^DNM1,2 DKO^ cells transiently expressing the main viral receptor ACE2^53^, and an EGFP-tagged version of the early endosome protein Rab5^54^, were used in these studies. Time-course uptake assays followed by confocal fluorescence microscopy were performed to distinguish the fraction of Alexafluor-555-labelled S localized at the PM from that internalized into endosomal vesicles labelled by Rab5-EGFP (early endosomes) (Figure 5). Tf labelled with Alexafluor-647 was used as an internal, positive control to monitor the extent of inhibition of dynamin-dependent endocytosis in each cell analysed. In vehicle-treated MEF^DNM1,2 DKO^ control cells, the uptake of S was efficient, the internalized protein colocalized with Rab5-EGFP positive vesicles (Figure 5 A-B, Ctrl). In dyn1,2 depleted MEF^DNM1,2 DKO^ cells most of the S fluorescent signal remained associated with the PM of the cells, even after 3h, indicating a strong inhibition of endocytosis (Figure 5 A-B, 4OH-TMX). Consequently, the colocalization of the intracellular Rab5 compartment with the S protein remained low throughout the time course (Figure 5 A-B, 4OH-TMX). Similar results were obtained for fluorescently labelled Tf, which also accumulated at the PM in dyn1,2 depleted MEF^DNM1,2 DKO^ cells at the expenses of intracellular compartments (Figure 5 A, 4OH-TMX). Taken together, these results demonstrate that, similarly to Tf, the ACE2-mediated endocytosis of SARS-CoV-2 S trimeric proteins is mainly dynamin-dependent.

**Figure 5.**
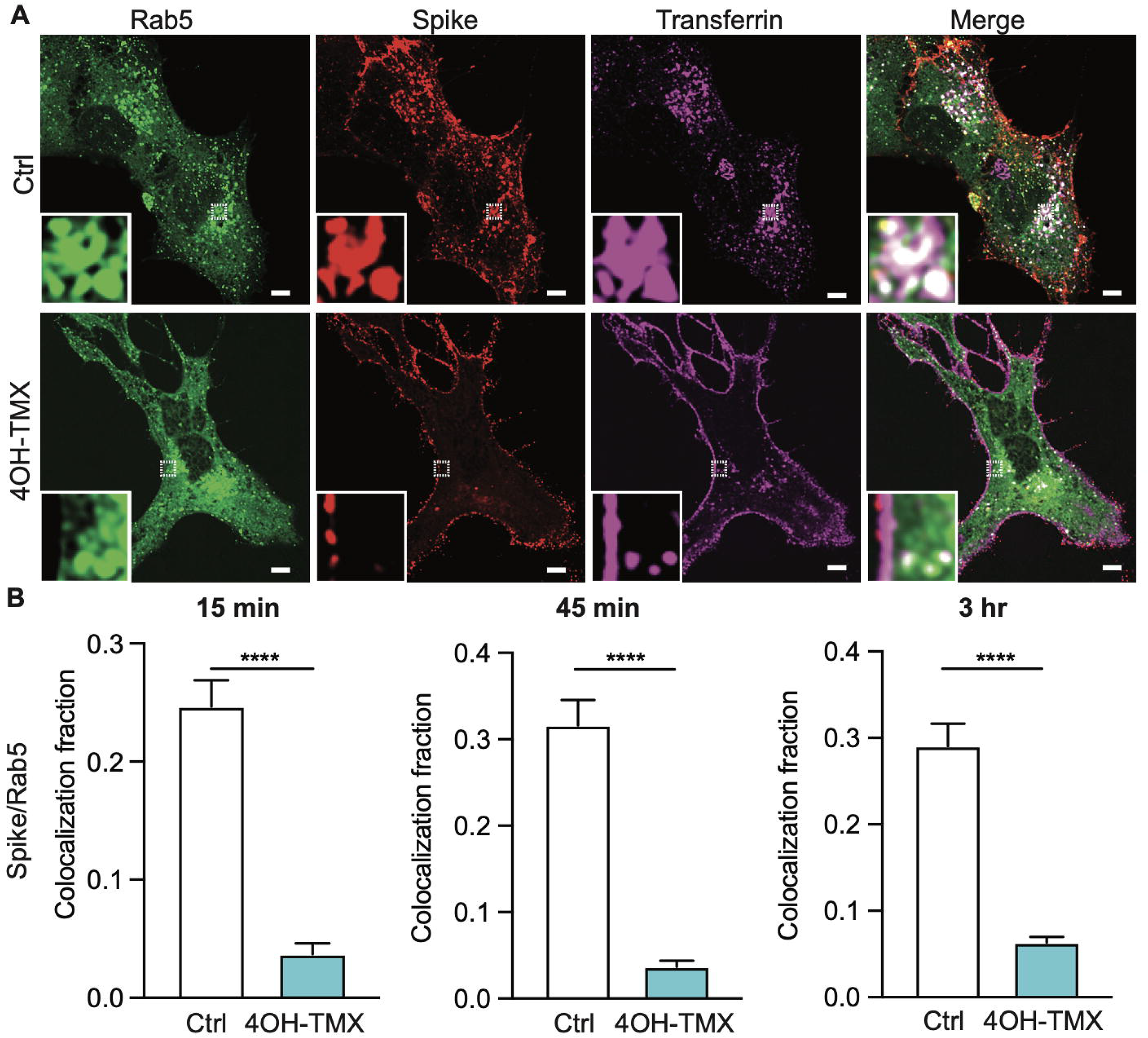
ACE2-mediated endocytosis of SARS-CoV-2 trimeric spike proteins is dynamin-dependent. A) Representative confocal fluorescence images of MEF^DNM1,2 DKO^ cells transiently expressing hACE2 and Rab5-EGFP, and treated with vehicle control or 4OH-TMX for 6 days and incubated for 3h with Alexa Fluor-555-labelled transferrin (Tf, magenta) and soluble trimeric SARS-CoV-2 Spike protein (red). The inset shows magnified images from the indicated white boxed areas. The images represent a single optical slice of the imaged cell. B) Quantification of colocalization between indicated proteins in cells treated as described in A). The mean±STDEV of 16 Ctrl cells and 14 4OH-TMX cells for 15 min; 16 Ctrl cells and 15 4OH-TMX cells for 45 min; and 17 Ctrl cells and 18 4OH-TMX cells for 3 h are shown. Statistical significance was calculated using a no parametric Mann-Whitney U test (****P<0.0001).

### SARS-CoV-2 infection in ACE2-expressing MEF cells requires endosome maturation and is insensitive to TMPRSS2 inhibitors

In addition to the MEF^DNM1,2, DKO^ cells, a triple dynamin 1,2,3 conditional KO cell line (here referred to as MEF^DNM1,2,3 TKO^)^55^ became available for this study. The dyn1,2,3-depletion in MEF^DNM1,2,3 TKO^ cells following a 6-day treatment with 4OH-TMX, blocks Tf uptake similarly to the dyn1,2-depletion in MEF^DNM1,2,3 DKO^ ^55^. An advantage of this triple dyn1,2,3 KO model system is that in addition to depletion of all three dynamin isoforms 1,2,3, these cells adhere better to culture surfaces compared to the MEF^DNM1,2, DKO^ cells. In addition, the responsiveness to 4OH-TMX treatment is more stable over cell passaging than in MEF^DNM1,2, DKO^ cells (not shown).

To study how a clinical isolate of the ancestral SARS-CoV-2 Wuhan strain (here referred to as Wuh^56^) infects MEF^DNM1,2,3 TKO^ cells, we created two transgenic cell lines, one stably expressing the human ACE2 gene (here referred to as MEF^DNM1,2,3 TKO^-ACE2), the other expressing the ACE2 gene together with a N-terminally GFP-tagged version of TMPRSS2 (here referred to as MEF^DNM1,2,3 TKO^-ACE2-TMP2-GFP). We used fluorescence activated cell sorting to isolate cells that expressed high, medium, and low levels of TMPRSS2-GFP. However, only the cells that expressed the lowels levels of GFP reattached to the culture flasks, indicating that in MEF cells the surface expression of the trypsin-like serine protease TMPRSS2 interferes with cell adhesion.

We first determined to which extent in these two cell lines the cell entry of SARS-CoV-2 Wuh strain depended on endosomal proteases or on the cell surface TMPRSS2. To this end, we tested the sensitivity of Wuh infection to nafamostat, an inhibitor of TMPRSS2^57^, and apilimod, an inhibitor of the phosphoinositol-5 kinase (PIP5K) required for efficient early to late endosome maturation and, therefore, delivery of endocytosed cargo to the lysosomal compartment^58,59^. Drug or vehicle-control pretreated infected cells were fixed at 20 hpi and the percentage of Wuh infection was monitored by immunofluorescence using antibodies against the viral protein N, followed by automated high-content imaging and image analysis (Figure S5 A). As expected, in cells that did not over-express TMPRSS2, Wuh infection was strongly inhibited by apilimod and not by Nafamostat (Figure S5 B and C, MEF ACE2). The over-expression of TMPRSS2-GFP, even if at low levels, had two main effects: Firstly, it slightly increased the overall infectivity of the virus in DMSO-control treated cells compared to the values obtained in cells that only expressed ACE2 (Figure S5 C, MEF ACE2 TMP2-GFP). Secondly, it rendered infection partially resistant to apilimod and sensitive to nafamostat (Figure S5 B and C). These results indicate that the MEF ACE2 cells either do not express TMPRSS2 or, if they do, the protein is not available to the virus. Hence, infection depends on virus endocytosis and delivery to endo/lysososmes. In MEF ACE2 TMP2-GFP cells, the low levels of TMPRSS2 at the cell surface provide the virus with access to an alternative entry pathway that bypasses endosome maturation.

### SARS-CoV-2 uses dynamin-dependent and -independent entry to infect ACE2-expressing MEF cells

The block of SARS-CoV-2 soluble trimeric spike S internalization observed in dynamin-depleted cells, and the sensitivity of infection to apilimod, indicated that SARS-CoV-2 infection of ACE2-expressing MEF cells requires endocytosis. To confirm that the results obtained with the S protein reflected the infectious entry pathway of the virus, infection assays were first performed in vehicle control and 4OH-TMX treated MEF^DNM1,2,3 TKO^-ACE2 cells. Two clinical isolates of SARS-CoV-2 were tested, the ancestral Whu^56^ virus and the more infectious Delta variant ^60^ (here referred to as Delta). In these experiments, we also investigated the role of the actin cytoskeleton using the actin depolymerising drug Lat-B. Because prolonged treatments with this drug result in loss of fibroblasts cell morphology and detachment, we implemented a procedure to interfere with actin dynamics only during virus entry, without compromising cell attachment (Figure 6 A). Six days after 4OH-TMX or vehicle control treatments, MEF^DNM1,2,3 TKO^-ACE2 cells were treated with 3 μM Lat-B for 15 min before infection. Lat-B was also present in the virus inoculum for 2 additional hours. Unbound virus and Lat-B containing medium was then removed and replenished with new medium containing 2 μM apilimod, to limit further infection after Lat-B removal (Figure 6 A). The fraction of infected cells was determined by immunofluorescence imaging and image analysis at 20 hpi (Figurre 6 B). In MEF^DNM1,2,3 TKO^ cells treated with 4OH-TMX for 6 days and DMSO-control, infection with Whu or Delta SARS-CoV-2 was reduced by up to 80% (Figure 6 C-D), indicating that in these cells, dynamin-dependent endocytosis represents the main productive viral entry route. Actin depolymerization inhibited infection by more than 60% in control cells and this inhibitory effect was even stronger in dynamin depleted cells for both tested viruses Whu and Delta (Figure 6 B-D). Thus, in ACE2-expressing MEF^DNM1,2,3 TKO^ cells, where TMPRSS2 is ether not expressed or not accessible to the virus, the infection of SARS-CoV-2 is mainly dynamin-dependent, and it is facilitated by the actin cytoskeleton. A similar dynamin- and actin-dependent entry mechanism has been described for other viruses, such as VSV^36^, that have a similar size to coronaviruses (i.e. 100-120 nm in diameter), and may not completely fit into dynamin-accessible endocytic invaginations. Actin polymerization facilitates the maturation of the endocytic cup, allowing the formation of the narrow membranous neck where dynamin binds and cleaves off the nascent vesicle^36^. Interestingly, similarly to other tested viruses but to a lower extent, increasing the virus load of SARS-CoV-2 Wuh to an amount sufficient to infect 25% of the cells restored infection up to 12% in a virus-dose dependent manner (Figure 6 E). Considering that 3-5% of these cells are not responsive to 4OH-TMX-induced depletion of dynamins, the remaining infection observed in dyn1,2,3-depleted cells indicated virus infection via dynamin-independent endocytosis. Thus, albeit with much lower efficiency compared to the dynamin-dependent entry, SARS-CoV-2 can also enter cells by an alternative endocytic mechanism.

**Figure 6.**
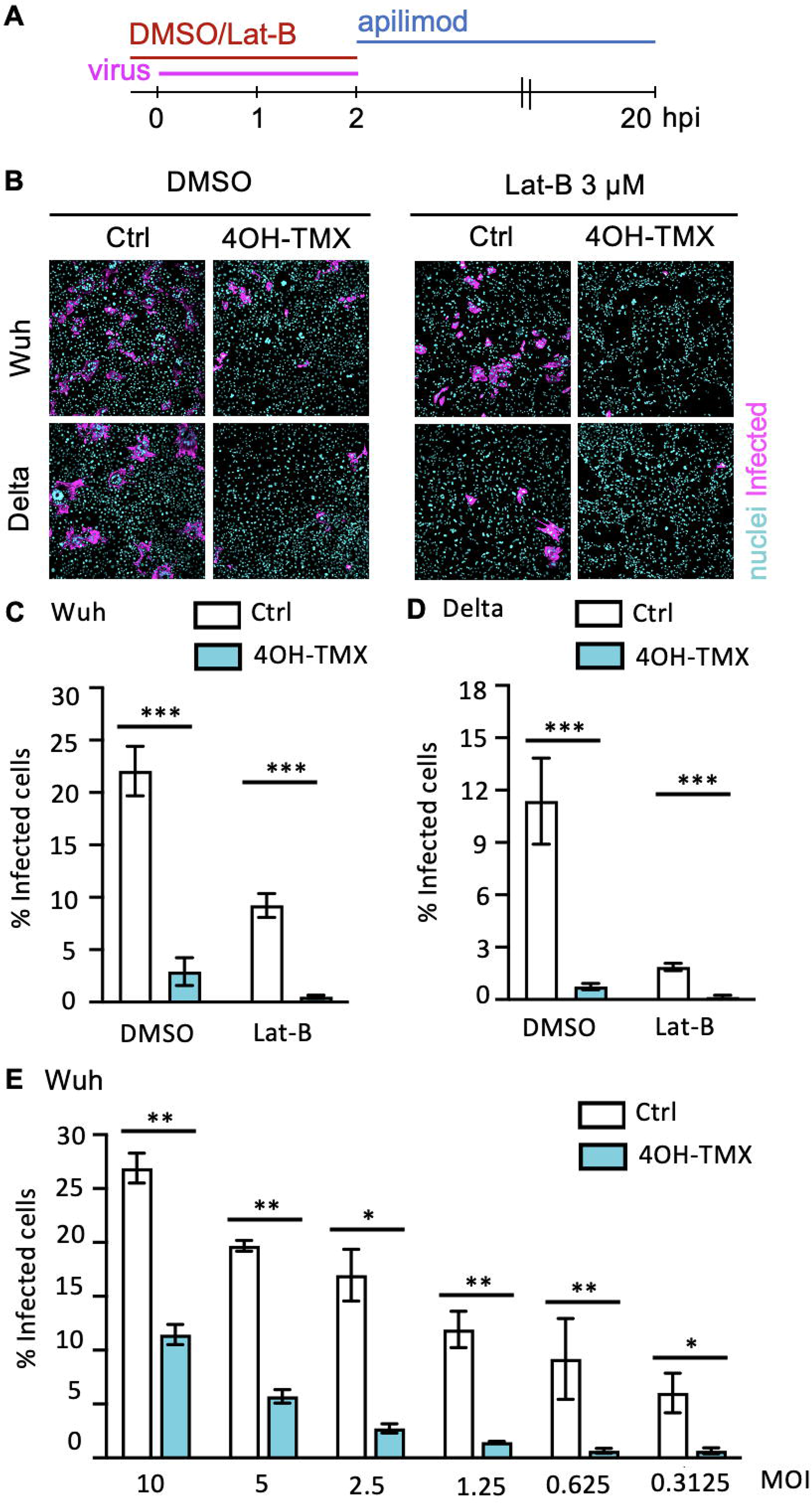
Dynamin-dependent and -independent entry mediate SARS-CoV-2 Wuh and Delta infection in ACE2-expressing MEF cells. A) Schematic description of the Lat-B treatment in MEF^DNM1,2,3 TKO^-ACE2. B) Representative fluorescence images of MEF^DNM1,2,3 TKO^-ACE2 treated with indicated compounds 15 minutes before infection and infected with Wuh or the Delta variant of SARS-CoV-2 for 20 h at 6 days after vehicle control (Ctrl) or 4OH-TMX treatment. C-D) Quantification by image analysis of SARS-COV-2 Whu or Delta infection in MEF^DNM1,2,3 TKO^-ACE2 cells treated with vehicle control (Ctrl) or 4OH-TMX for 6 days and infected with indicated MOIs for 20 h. Values indicate the mean of at least three independent experiments and the error bars represent the STDEV.(E) Quantification by image analysis of SARS-COV-2 Wuh infection in MEF^DNM1,2,3 TKO^-ACE2 cells treated with vehicle control (Ctrl) or 4OH-TMX for 6 days and infected with indicated MOIs for 20 h. Statistical significance was calculated using a no parametric Mann-Whitney U test (*p<0,05; **p<0,01; ***p<0,001).

### Depletion of dynamins blocks endocytosis of SARS-CoV-2 virions

To confirm that the inhibition of infection observed in dynamin depleted cells corresponded to a proportional block in endocytosis of SARS-CoV-2 virions, we directedly measured the extent of virus internalization in dyn1,2,3-depleted and controls MEF^DNM1,2,3 TKO^ -ACE2 cells. Viruses (Wuh, equivalent MOI of 50) were added to cells at 37°C for 60 min, in the presence of 50 μM cycloheximide to prevent viral protein synthesis. After fixation, cells were processed for sequential immunofluorescence staining to distinguish viral particles outside of the PM, i.e. particles not yet internalized (virus out), from internalized virions (virus in) (Figure 7 A). In this technique, non-internalized viruses are immunostained in fixed cells before permeabilization, using a combination of polyclonal antibodies against the viral spike protein, followed by secondary antibodies conjugated to a fluorophore (Figure 7A, Ab 1, magenta). After a second fixation, cells are permeabilized and stained again with the same anti-spike antibodies followed by secondary antibodies conjugated to a different fluorophore (Figure 7 A, Ab 2, green). Hence, in this assay, non internalized virions are stained either with the first (i.e. magenta spots) or with both (i.e. white colocalizing spots) fluorophores (Figure 7 B, Ctrl, magenta or white dots). Internalized virions are stained only with the second fluorophore (Figure 7 B, 4OH-TMX, green dots)

**Figure 7.**
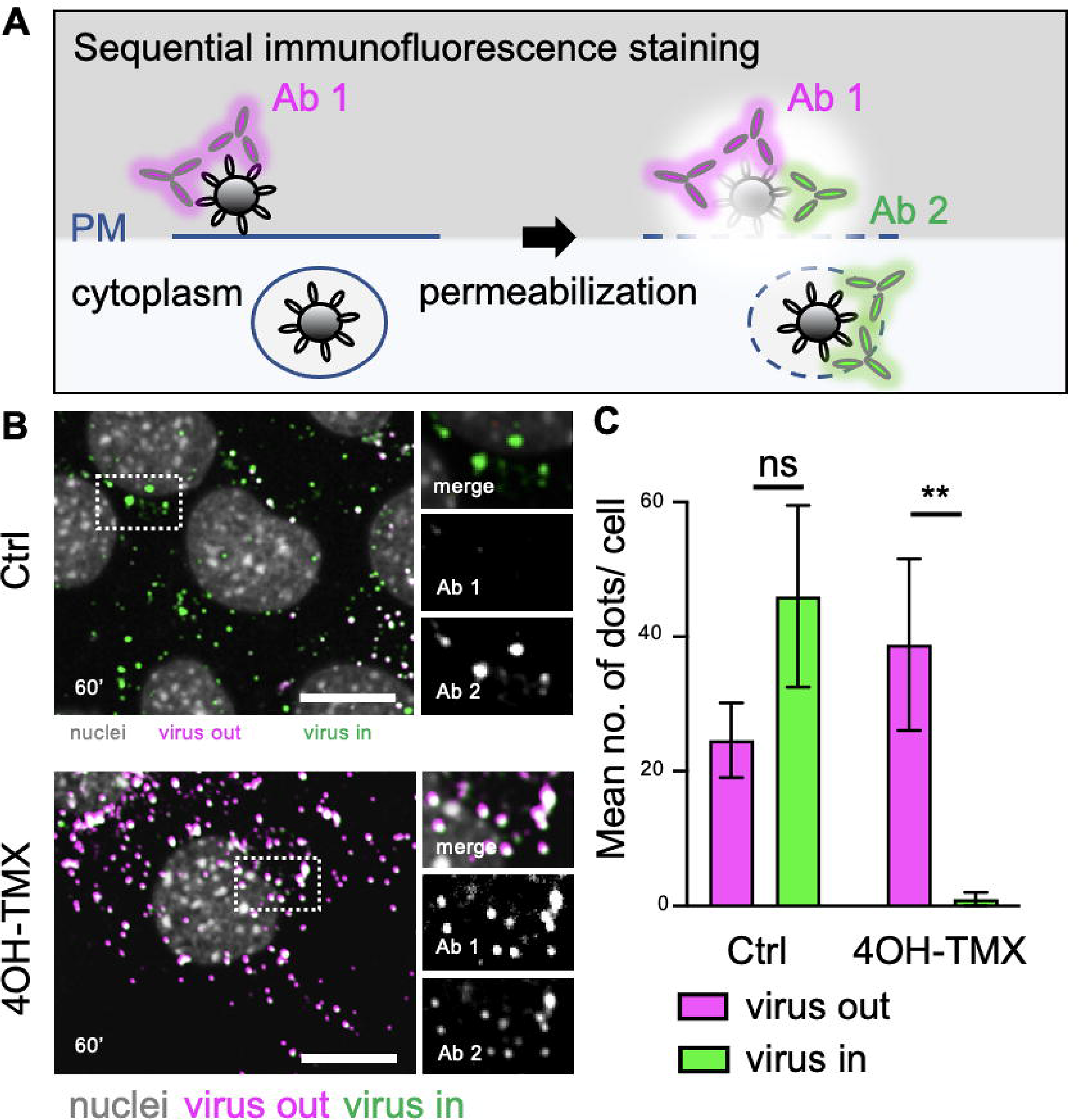
Depletion of dynamins 1-3 blocks SARS-CoV-2 endocytosis in ACE2 expressing MEF cells. A) Schematic representation of the sequential immunostaining protocol. After fixation, and before permeabilization viruses on the surface of MEF are immunostained before permeabilization using antibodies against the spike protein (Ab 1) followed by secondary antibodies conjugated to a fluorophore (e.g., excitation 488nm, virus out). After a second fixation and permeabilization, the immunostaining is repeated (Ab2) but using secondary antibodies conjugated to a different fluorophore (e.g., excitation 647nm, virus in). B) Representative images of virus entry in MEF^DNM1,2,3 TKO^-ACE2 cells (4OH-TMX) or control MEF cells (Ctrl) fixed at 60 min after virus inoculation and processed for sequential immunofluorescence as described in A. Inset images show a magnification of the area indicated by the white dashed boxes, with merged as well as separated fluorescence images. Virus out= non internalized viruses; virus in= internalized viruses. C) Quantification of non-internalized (virus out) and internalized (virus in) virions using automated image analysis. Values represent the mean of 15 cells from 3 independent experiments, and error bars represent the STDEV. Scale bars= 10 μm Statistical significance was calculated using XXX (**p<0,01; n.s.=non-significant).

In the unperturbed control cells, at 60 min post infection, image analysis after confocal imaging revealed that a sizable fraction of viruses was found inside the cells (Figure 7 C, Ctrl). In contrast, depletion of dyn1,2,3 blocked virus endocytosis almost completely (Figure 7 C, 4OH-TMX). This experiment demonstrates that the strong block of virus infection observed in dyn1,2,3-depleted cells coincides with a proportional block in virus endocytosis.

### TMPRSS2 expression bypasses the need for dynamin-dependent endocytosis

As opposed to small-molecule inhibitors, RNAi or DN approaches, which suffer of uncharacterized off-targets effects or incomplete inactivation of dynamins, the results presented here were obtained in cells where all three isoforms of dynamins were completely depleted by a chemically induced KO. Although coronaviruses are potentially able to infect cells by direct fusion with the PM, the exact mechanism of virus infection, i.e. the extent of endocytosis-vs PM-fusion, in different cell types *in vivo* is not fully understood. To determine whether low levels of TMPRSS2 could rescue infection in dynamin-depleted cells, infection assays were repeated in vehicle control and 4OH-TMX treated MEF^DNM1,2,3 TKO^-ACE2-TMPRSS2-GFP cells. Six days after 4OH-TMX or vehicle control treatment, cells were infected with SARS-CoV-2 Wuh for 20 h and the fraction of infected cells was determined by immunofluorescence imaging and image analysis at 20 hpi. As opposed to a strong reduction of infection in cells that do not express TMPRSS2 (Figure 8 A-B, MEF ACE2), and despite the low levels of TMPRSS2-GFP expression that we could isolate by FACS, the presence of this serine protease rescued infection in dynamins-depleted cells up to 50% of the levels obtained in vehicle-control treated cells (Figure 8 A-B, MEF ACE2 TMP2-GFP). Thus, TMPRSS2 allows infection in a dynamin-independent manner. Whether this infection is mediated by direct fusion of viruses at the PM or by dynamin-independent endocytosis remains to be established.

**Figure 8.**
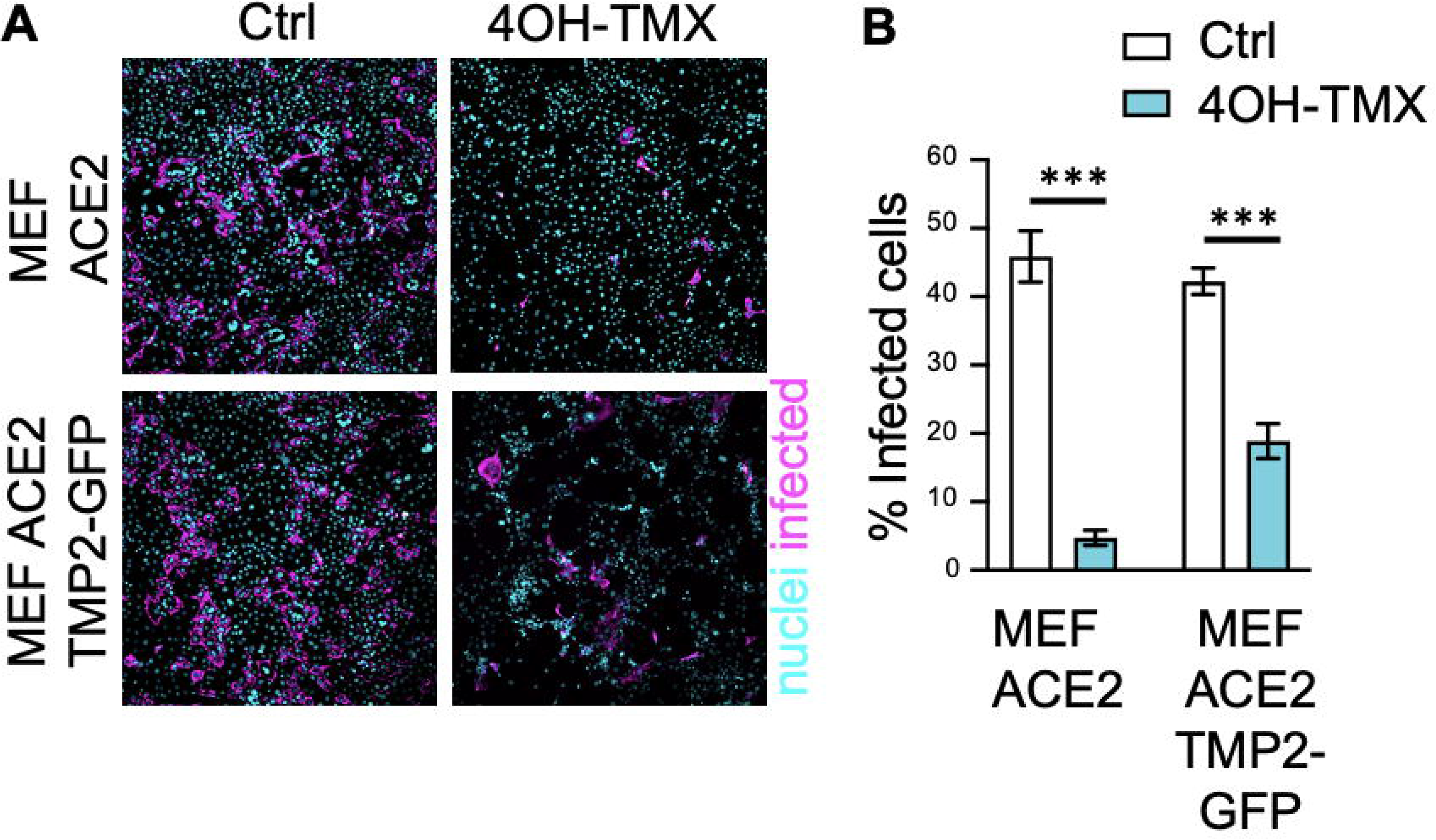
TMPRSS2 expression bypasses the need for dynamin-dependent endocytosis. A) Representative fluorescence images of MEF^DNM1,2,3 TKO^ ACE2 and MEF^DNM1,2,3 TKO^ ACE2-TMPRSS2-GFP treated with EtOH vehicle control (Ctrl) or 4OH-TMX for 6 days and infected with SARS-CoV-2 for 20 hours. Infected cells were identified after immunostaining using antibodies against the viral NP protein (magenta). Nuclei stained with the DNA dye Hoechst (cyan). B) Quantification by image analysis of SARS-CoV-2 Wuh infection in MEF^DNM1,2,3 TKO^-ACE2 and MEF^DNM1,2,3 TKO^ ACE2-TMPRSS2-GFP cells treated with EtOH vehicle control (Ctrl) or 4OH-TMX for 6 days and infected with indicated for 20 h. Values indicate the mean of at least three independent experiments and the error bars represent STDEV. Statistical analysis was performed using an unpaired double tailed t-test (*** p<0.001).

## Conclusions

While inhibiting one endocytic pathway results in the block of internalization for receptors such as TfR and its ligands Tf and CPV, many viruses can enter cells and initiate their infection cycle using more than one endocytic mechanism. For viruses whose infection was affected by dynamin depletion, such as SARS-CoV-2, influenza and alphaviruses, the inhibitory effect was more evident at low MOI. Increasing the dose of virus per cell gradually restored infection, for viruses such as SFV completely. This could be interpreted at least in two ways: i) dynamin depletion blocks internalization of the main viral receptor, and the dynamin-independent mechanism is mediated by either alternative, less accessible receptor(s), or by unspecific binding to charged molecules at the cell surface that are internalized via a dynamin-independent pathway. In support of this scenario, previous work has shown that over-expression of temperature sensitive dynamin mutants, which at the restrictive temperature acted as dominant negatives, increased the dynamin-independent uptake of fluid phase markers^61^. Similarly, in the dynamin depleted MEF cells used here, fluid phase uptake is increased^19,55^. ii) The main virial receptor could be internalized using two endocytic mechanisms, one mainly available in unperturbed cells (e.g. dynamin-dependent), the other activated upon dynamin depletion (i.e. dynamin independent). In the case of SARS-CoV-2, this second scenario is plausible because in the absence of ACE2 expression, dyn1,2,3 depleted cells were not infected (not shown), indicating that the dynamin-independent entry still requires the viral receptor ACE2. Whether other SARS-CoV-2 host entry factors, such as the cell co-receptor neuropilin-1^56^, which is internalized via dynamin-independent endocytosis^62,63^, could contribute to dynamin-independent virus entry will require further studies.

Currently, the dominant SARS-CoV-2 variant is omicron. Recent evidence has shown that this variant favours endosomal entry rather than direct fusion at the PM^16^. The implication of these findings is that variants such as omicron, which still causes thousands of deaths worldwide every month, would strongly relay on endocytosis to access the endo/lysosomal compartment. Based on our results, we propose that dynamin inhibitors could have beneficial therapeutic potential in combination with other antivirals.

## Acknowledgments

We thank the University of Helsinki Graduate Program in Microbiology and Biotechnology for supporting R.O. The Academy of Finland Research Grants 335527 (G.B), the Jane and Aatos Erkko Foundation (O.P.V.) Helsinki University Hospital funds TYH2021343 (O.P.V.), European Union’s Horizon Europe Research and Innovation Program grant 101057553 (G.B., O.P.V.), The University of Queensland Amplify Fellowship (M.J.), the Australian National Health and Medical Research council (grant 2010917 G.B. and F.A.M.). This work was also supported by Australian Research Council (ARC) Discovery Early Career Researcher Award (DE190100565) and The University of Queensland Amplify fellowship to M.J.. The work was supported by an ARC Linkage Infrastructure Equipment and Facilities grant (LE130100078) and an National Health and Medical Research Council (NHMRC) Senior Research Fellowship (GNT1155794) to F.A.M. P.Y.L is supported by the Agence Nationale de la Recherche (ANR) funding (grant numbers ANR-21-CE11-0012 and ANR-22-CE15-0034). Work in the Greber lab was supported by grants from Schweizerischer Nationalfonds (Swiss National Science Foundation 31003A_179256 and 310030_212802). We are grateful to the Jane and Aatos Erkko Foundation for support to M.V.R. and the Academy of Finland under award numbers 330896 to M.V.R.) and 332615 to E.M. J.M. was supported by core funding to MRC Laboratory for Molecular Cell Biology at University College London (MC_UU_00012/7).

High-throughput imaging was performed at the Light Microscopy Unit of the University of Helsinki.

The funders had no role in the study design, data collection, or interpretation. We declare no competing interests.

## Materials and Methods

### Cells

MEF cells monolayers were grown in high-glucose Dulbecco’s Modified Eagle Medium (DMEM; Merk, cat. no. D6546) containing 100 IU/ml of streptomycin and penicillin (Merk, cat. no. P0781), and 10% heat inactivated fetal bovine serum (FBS; Thermo Fisher Scientific, cat. no. 10270-106) in a 37°C incubator with 5% CO_2_. MEF^DNM1,2 DKO^ and MEF^Dyn1,2,3 TKO^ cells were a kind gift of Dr. Shawn Ferguson and Pietro De Camilli^19,55^. The depletion of dynamins in these cells was induced by incubating cells with 3 μM 4OH-TMX (dissolved in EtOH) for two consecutive days. The medium was then removed, and cells grown in new medium containing 0.3 μM 4OH-TMX. At day 5 post treatment, the medium was removed, and cells seeded in 96-well imaging plates (PerkinElmer, cat. no. 6005182), at a density of 15,000 (vehicle) or 30,000 (4OH-TMX) cells per well. To obtain equal cell densities in the study, seeding a double number of dynamin-depleted cells was necessary because complete dynamin depletion blocks cell division after day 5 post 4OH-TMX induction for colocalization of soluble trimeric spike with Rab5-EGFP, cells were seeded on fibronectin-coated glass coverslips in 24 well plates at a dilution of 50,000 (vehicle) or 100,000 (4OH-TMX) cells per well. Experiments were performed at day 6 post 4OH-TMX treatment.

To generate MEF^DNM1,2 DKO^ transiently expressing human ACE2 and Rab5-EGFP, cells were electroporated with a Biorad Gene Pulser Xcell system, using PBS and 2 μg of plasmid DNA (plenti-6.3-hACE2^56^, pRab5-EGFP^54^). For CPV infections, MEF cells was transfected with feline TfR1 plasmid (pCTfR, 2.5 μg)^25^ by using TransIT-LT1transfection reagent (Mirus Bio, WI, USA). To generate MEF^Dyn1,2,3 TKO^ cells stably expressing hACE2, cells were transduced with the respective lentivirus (pWPI-IRES-Puro-Ak-ACE2, kindly provided by Dr. Sonja Best, Addgene viral prep # 154985-LV) at MOI 0.3. At 48 hpi, positively transduced cells where selected with puromycin (1 μg/ml) for 5 days and the resistant cells amplified and stored in liquid nitrogen.

To generate MEF-ACE2 expressing TMPRSS2-EmGFP cells were seeded (1×10^6^/well) in a 6-well plate and grown overnight in DMEM media with 10% FBS, 2 mM l-glutamine, 1% penicillin-streptomycin. Each well received 200 µl of a solution containing 3×10^5^ infectious units of a commercially available third-generation lentivirus expressing the gene of human *TMPRSS2* with a C-terminally fused histidine tag followed by the mGFP protein, and from a separate promoter the puromycin resistant gene (Angio-Proteomie, catalogue number vAP-0101). Lentiviral infections were carried in Dulbecco’s modified eagle’s medium (DMEM), 2 mM glutamine, 0,5 % bovine serum albumin (BSA), and 1x penicillin/streptomycin antibiotic mix, at an MOI of 0.3 infectious units/cells (the lentivirus titre was provided by the manufacturers). After selection for 7 days with 3 mg/ml puromycin, cells were amplified for three passages and then then FACS sorted in high, medium, and low TMPRSS2 expressing cells. Only the low TMPRSS2-mGFP expressing cells survived in culture, and were used in the study.

### Viruses

All experiments with wild-type or mutant SARS-CoV-2 were performed in BSL3 facilities of the University of Helsinki with appropriate institutional permits. Viruses isolated from COVID-19 patients were propagated once in VeroE6-TMPRSS2 cells, titrated by plaque assay, and stored at −80°C in DMEM, 2% FBS, P/S. All SARS-CoV-2 viruses were sequenced by next generation sequencing at the University of Helsinki. SFV-EGFP^64^, SINV-mCherry^65^, VSV-EGFP^37^ and UKNV^66^ have been described and were propagated for 22 h in BHK 21 cells in DMEM, non-essential amino acids, 1xGlutamax, 1x Penicyllin/Strempomycin, and 20 mM HEPES pH 7.2. VACV-EGFP^28^ was propagated in Hela cells. Influenza A X31 strain was propagated in A549 cells^67^. CPV type 2 was grown, isolated and concentrated as previously described^25^. AdV-C5_EGFP is an E1/E3 deletion mutant virus with E1 region replaced by the enhanced green fluorescent protein (EGFP) gene under the control of cytomegalovirus major immediate early promoter^39^. The virus was grown in 911 cells and purified on CsCl gradients as previously described^68,69^. HRVA1A stock virus was produced as previously described^67^.

### Small molecule inhibitors and protein-labelling reagents

All small molecule inhibitors were purchased from Tocris except for Lat-A, Lat-B and 4OH-TMX (Sigma). Drugs were solubilized in DMSO and stored at −20°C, were used at the following concentration unless otherwise indicated in the result session: DMSO 0.1%, Bafilomycin A1 50 nM, NH_4_Cl 20 mM, sucrose 0.45 mM, Chlorpromazine 20 mM, Pitstop 25 μM, Dyngo-4a 30 μM, Dynole 30 μM, Dynasore 80 μM, Cytochalasin-D 5μM, Lat-A 3 μM and Lat-B 1 μM, LatA 0.1 µM, EIPA 80 μM, Wortmannin 200 nM, ML142 10 μM. Tf-A647 (5 μg/ml, Thermo Fisher Scientific, cat. no. T23366) and CTB-A647 (Thermo Fisher Scientific, cat. n. C34778). For actin filament labelling, Alexa Fluor 647 Phalloidin was used (ThermoFisher, cat. no. A22287). To fluorescently label the soluble trimeric SARS-CoV-2 spike we used Atto-550 NHS-ester labelling reagent, (Atto-Tec GmbH, cat.n. AD550-35) according to manufacturer instructions for one hour at room temperature in the dark. The remove the excess unbound fluorophore the mixture (500 ml) was passed through NAP-25 Sephadex (GA Healthcare) columns and eluted in 750 ml of PBS according to manufacturer instructions.

### Fluorescence-activated cell sorting (FACS) analysis

After infection, cells in 24 well plates were washed once in PBS and detached by incubation in 300 μl of 0.25% Trypsin/EDTA solution for 10 min at room temperature. Trypsin was blocked by addition of 300 μl of DMEM containing 10% FBS and cells fixed by addition of 600 μl of 8% PFA. Fixed cells were incubated for 20 min at room temperature in a rotator. The fixation reaction was blocked by addition of NH_4_Cl to a final concentration of 20 mM. Cells were centrifuged at 1000 g for 10 min, and the pellet resuspended in PBS. In the case of viruses that expressed fluorescent reporter proteins (i.e. SFV-EGFP, VSV-EGFP, VACV-EGFP, and SIN-mCherry), infected cells were directedly analysed by FACS. In the case of CPV, IAV and UUKV virus, cells were permeabilized with 1% saponin (freshly prepared) in PBS containing 0.5% BSA and incubated with primary and secondary antibodies in the presence of 0.5% saponin in PBS containing 0.5% BSA. Antibodies to detect CPV NS1^25^, IAV NP^67^, and Uukuniemi virus E^66^ proteins have been described. Samples were analysed for viral antigens fluorescence (or viral induced EGFP or mCherry) by a FACS ARIA (BD Biosciences, NJ, USA) and FACSCalibur flow cytometers (BD Biosciences) systems. Non infected, drug treated cells were used to set the minimal threshold fluorescence levels (the ‘gates’) during FACS analysis.

### AdV-C5 and HRV-A1 infections and immunofluorescence in MEF^DNM1,2 DKO^

MEF^DNM1,2, DKO^ cells were seeded into 25 cm^2^ flask in growth medium containing 2 µM 4OH-tamoxifen (Sigma, cat. no. H-6278) for 48 h. Medium was replaced by growth medium containing 300 nM tamoxifen and after two days, cells were trypsinized and seeded into 96-well imaging plates (Greiner Bio-One, cat. no. 655090). Cells were further incubated in medium containing 300 nM tamoxifen for 48 h and then incubated with two-fold dilutions of AdV-C5-EGFP stock virus in DMEM medium (Sigma-Aldrich, cat. no. D6429) supplemented with 7.5% fetal calf serum (Gibco/ Thermo Fisher Scientific, 10270106), 1% non-essential amino acids (Sigma-Aldrich, cat. no. M7145) and 1 % penicillin-streptomycin (Sigma-Aldrich, cat. no. P0781). After 22 h, cells were fixed with 3% PFA in phosphate-buffered saline (PBS) for 30 min at room temperature, quenched with 25 mM ammonium chloride in PBS for 10 min and nuclei were stained with 4’,6-diamidino-2-phenylindol (DAPI, Sigma, cat. no. D9542; 1 µg/ml solution in PBS/0.1% TritonX-100) for 20 min. The plate was imaged with a Molecular Devices automated ImageXpress Micro XLS as described^2^ using 10 × SFluor objective (NA 0.5). Images were analyzed using CellProfiler^70^ and mean EGFP intensity over the DAPI mask was determined. Non-infected control wells were used to determine the threshold for an infected cell, threshold being the mean of maximum values from non-infected cells. Knime Analytics Platform (https://www.knime.com/knime-analytics-platform) was used to calculate infection indexes, i.e., fraction of EGFP-positive cells over total number of cells analyzed. HRV-A1a infection assay was carried out by incubating cells with two-fold dilutions of HRVA1A stock virus in plain DMEM at 37°C for 24 h. Subsequently, cells were fixed and processed for immunostaining with mouse anti-VP2 (RT16.7) and secondary goat Alexa Fluor 488-conjugated anti-mouse antibodies (Thermo Fisher Scientific, cat. no. A-11029) as described^2,71^. The plate was imaged with a Molecular Devices automated ImageXpress Micro XLS with 20 × SFluor objective (NA 0.75). Images were analyzed using CellProfiler and mean antibody signals were detected over a cell area corresponding to the DAPI mask extended by three pixels. Non-infected control wells were used to determine the threshold for an infected cell, threshold being the 90% cut-off value from the noninfected cells. Knime Analytics Platform was used to calculate the infection indexes.

### Immunofluorescence assay for CPV, UUKV and IAV

Cells fixed in 4% PFA for 20 min at room temperature (RT), were then incubated with a blocking solution of PBS containing 20 mM NH_4_Cl, for 20 min and then permeabilized with 0.1% Triton X-100 in PBS for 10 minutes at RT. After two washes with Dulbecco PBS, cells were incubated with primary antibodies against viral proteins for 2 h at room temperature. After two washes in Dulbecco PBS, cells were incubated with fluorescently conjugated secondary antibodies containing 1μg/ml DAPI for 1 h at RT. After three washes in PBS, cells in 96-well imaging plates were processed for imaging. Cells in coverslips were rinsed with milli-Q water and mounted on glass slides using Prolong Gold anti fade mounting medium (Thermo Fisher Scientific, P101144) and stored at 4°C in the dark until imaging.

### Confocal imaging and colocalization analysis

Samples on coverslips (i.e. internalization of SARS-CoV-2 soluble trimeric spike) were imaged with a W1 Yokogawa Spinning Disk confocal microscope using a 60x oil immersion objective. Images were acquired with a Hamamatsu ORCA-Flash4.0 V2 sCMOS (photon conversion factor 0.46) camera. Colocalization was performed with Imaris software to quantify the fraction of internalized spike particles colocalizing with the endocytic markers Rab5-EGFP and Tf-A647. For each three-dimensional confocal stack, spike particles and endocytic vesicles were automatically detected using the Imaris spot colocalization tool as described^26^.

### Virus entry by sequential immunostaining

The extent of virus internalization in dynamin-depleted and control MEF^Dyn1,2,3 TKO^-ACE2 cells was determined by immunofluorescence staining followed by confocal imaging and image analysis. Viruses (equivalent MOI of 50) were added to cells in 96 well imaging plates (PerkinElmer, CellCarrier 96 ultra, 6055302) at 37°C for 60 min, in the presence of 50 μM cycloheximide to prevent viral protein synthesis. After fixation, cells were processed for sequential immunofluorescence staining to distinguish viral particles on the outer leaflet of the plasma membrane, i.e. particles not yet internalized, from internalized virions. Non-internalized viruses were immunostained in fixed cells (4% PFA, 10 min, room temperature) before permeabilization, using a combination of polyclonal antibodies against the viral spike protein (1:200 dilution of anti S1 RBD cat. no. 40592-T62, and 1:200 dilution of anti S2 cat. no. 40590-T62, both from Sino Biological), for 3 h at room temperature. After 3 washed with PBS supplemented with 0.5% BSA (PBS-BSA), cells were incubated with secondary antibodies conjugated to an Alexa-488 fluorophore (1:500 dilution of goat anti rabbit Alexa Fluor-488; Thermo Fisher Scientific, cat. no. A32731), for 2 h at room temperature. After 2 washes with PBS-BSA, and a second fixation with 4% PFA for 5 min at room temperature, cells were washed 3 times with PBS-BSA, and permeabilized with 0.1% Triton-X 100 for 10 min at room temperature. After 3 washes in PBS-BSA, cells were stained again with the same anti-spike antibodies (dilution 1:200) for 3 h at room temperature. After three washes in PBS-BSA, cells were incubated with DNA staining Hoechst (Thermo Fisher Scientific, cat. no. H3569) and secondary antibodies conjugated to a different fluorophore, Alexa-647 (1:500 dilution of goat anti rabbit Alexa 647; Thermo Fisher Scientific, cat. no. A32733), for 2 h at room temperature. After 3 washes in PBS, cells were imaged by spinning disk confocal microscopy using a followed by high content spinning disc confocal imaging using a Yokogawa CQ1 microscope equipped with a 40x air objective. Automated image analysis to detect particles stained by both fluorophores (i.e. particles at the cell surface) and particles stained only by the second fluorophore (i.e. particles inside the cell) was performed with the Yokogawa Cell Path Finder software.

### Single-molecule imaging and particle tracking

Time-lapse total internal reflection TIRF microscopy imaging for live MEF cells was conducted on TIRF microscope (Roper Technologies) equipped with an ILas2 double-laser illuminator (Roper Technologies), a CFI Apo TIRF 100× (1.49-NA) objective (Nikon), and two Evolve512 delta EMCCD cameras (Photometrics). Image acquisition was performed using Metamorph software (version 7.7.8; Molecular Devices). Cells were bathed in buffer A (145 mM NaCl, 5 mM KCl, 1.2 mM Na_2_HPO_4_, 10 mM D-glucose, and 20 mM HEPES, pH 7.4) at 37°C. Single molecule movies were captured at 50-Hz (16,000 frames by image streaming) and 20-ms exposure for both controls, 4OH-TMX treated and Lat-A treated MEF cells. A quadruple beam splitter (LF 405/488/561/635-A-000-ZHE; Semrock) and a QUAD band emitter (FF01-446/510/581/703-25; Semrock) were used to isolate the LifeAct-mEos2 signal from autofluorescence and background noise signals. Lat-A (0.1 µM) was added 15 minutes before starting the imaging acquisition to depolymerize the actin filamin. To photoactivate the LifeAct-mEos2 molecules, a 405-nm laser was used with a 561-nm laser simultaneously to excite the photoconverted molecules. PALMTracer ^72,73^ in Metamorph software (MetaMorph Microscopy Automation and Image Analysis Software, v7.7.8; Molecular Devices) was used to obtain the mean square displacement (MSD) and diffusion coefficient (D; μm^2^ s^−1^) values. Tracks shorter than eight frames were excluded from the analysis to minimize nonspecific background. The Log_10_D immobile and mobile fraction distributions were calculated as previously described ^72^, setting the displacement threshold to 0.03 μm^2^ s^−1^ (*i.e*. Log_10_D = −1.45 when [D] = μm^2^ s^−1^). The mobile to immobile (M/IMM) ratio was determined based on the frequency distribution of the diffusion coefficients (Log_10_D) of immobile (Log_10_D ≤ −1.45) and mobile (Log_10_D > −1.45) molecules (the immobile fraction of molecules represents LifeAct-mEos2 molecules for which the displacement within 4 frames was below the spatial detection limit of our methods, 106 nm). The area under the MSD curve (AUC) was calculated in Prism 9 for macOS version 9.1.1 (GraphPad Prism 9 for macOS; https://www.graphpad.com/scientific-software/prism/). The super-resolved image color-coding was done as previously described ^74^ using ImageJ/Fiji (2.0.0-rc-43/1.50e; National Institutes of Health), with each colored pixel in the average intensity maps indicating the localization of an individual molecule (bar: 8 to 0, high to low density), the color-coded pixels in the average diffusion coefficient map presenting an average value for each single-molecule track at the site of localization (bar: Log_10_ 1 to −5, high to low mobility), and the color-coding of the track maps representing the detection time point (bar: 0–16,000 frame acquisition) during acquisition.

### Western blotting

MEF cells, either control or 4OH-TMX treated, were harvested with same number of cells on day6 to day9 after 4OH-TMX (or EtOH as vehicle control) was introduced to the cells. MEF cell lysate was mixed with same volume of 2× Laemmli sample buffer, boiled for 5 min at 95°C, and run on 10%-15% SDS-PAGE gels. Wet Transfer was then conducted on ice with a PVDF transfer membrane (ImmobilonTM-P), following the Western blotting with the mouse anti-Dyn1,2 antibody (BD Bioscience, cat. no. 610245). β-actin was used as loading control to show the overall actin express level was not changed with 4OH-TMX treatment, with mouse anti–β-actin antibody (Abcam, cat. no. ab6267).

### Electron microscopy

SFV entry was studied using transmission electron microscopy (TEM) analysis in MEF^DNM1,2 DKO^ cells and control MEF cells. Cells were grown on fibronectin-coated glass coverslips and the cells were treated with 4OH-TMX or vehicle (EtOH) control for 6 days and, following virus adsorption at 4°C for 1 h, cells were shifted at 37°C for 10 minutes to allow virus entry. Coverslips were then fixed and processed for flat embedding as described earlier^75^. Thin sections (60 nm) were cut parallel to the coverslip and imaged with Jeol JEM-1400 microscope operated at 80 kV. Images were acquired with Gatan Orius SC 1000B camera from approximately same depth of the cells, capturing 120 images from both conditions each time a SFV was observed. The distribution (% of total) of SFV in CCPs (identified based on clathrin coated endocytic profile), in non-coated profiles (identified based on absence of clathrin coat) and other endocytic profiles (mainly large endocytic structures) was then counted manually and is presented as average.

---

**Figure S1.**
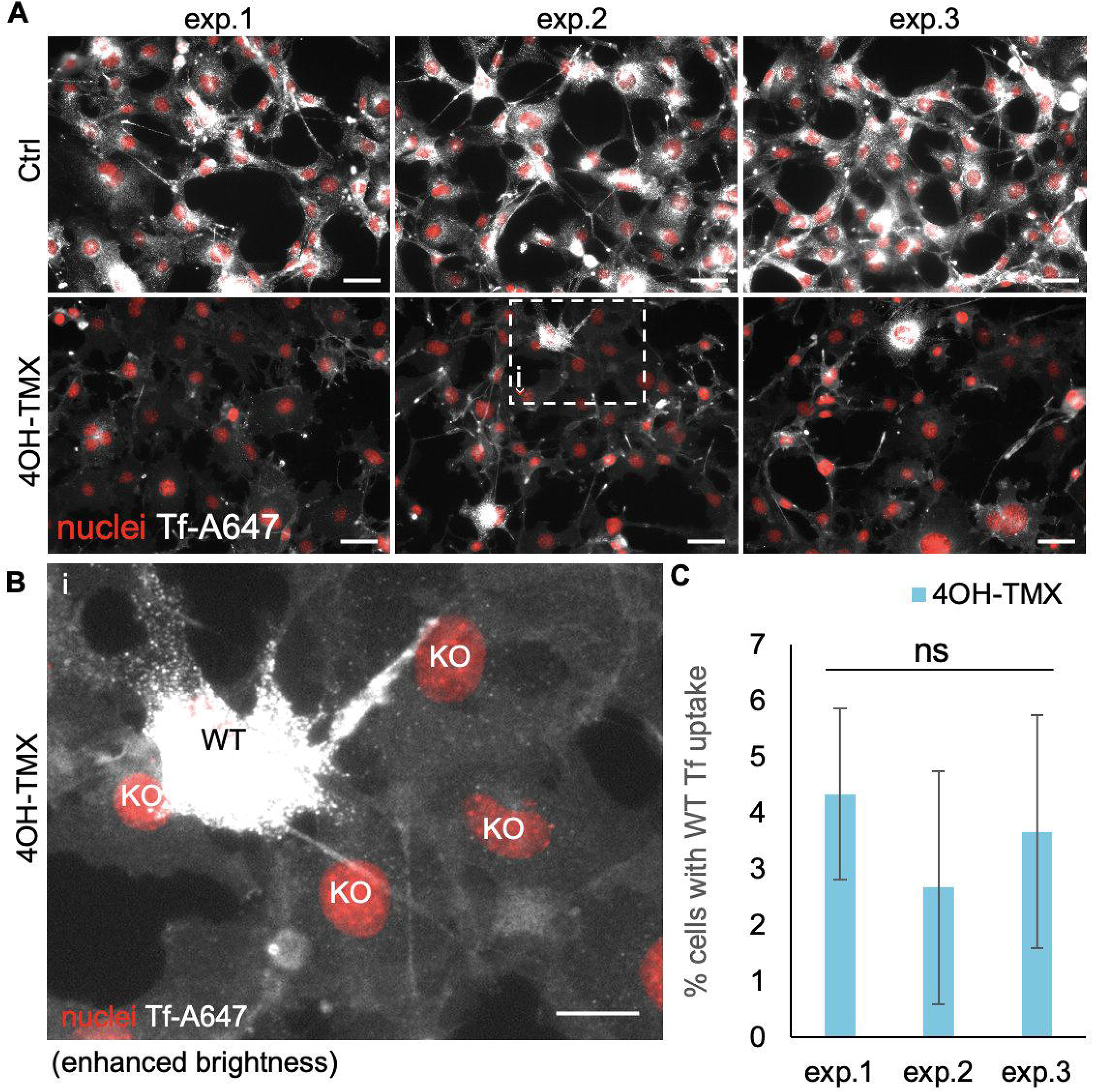
A small fraction of MEF^DNM1,2 DKO^ cells do not respond to 4OH-TMX treatment. A) Representative fluorescence images of three independent experiments where 20 min uptake of 5 mg/ml Tf-A647 at 37 °C was monitored in MEF^DNM1,2 DKO^ pre-treated with EtOH vehicle control (Ctrl) or 4OH-TMX for 6 days. Scale bar= 50 μm, nuclei stained with Hoechst (red). B) The white boxed area from panel A was expanded to distinguish cells where the uptake of Tf-A647 was blocked (KO; i.e. the 4OH-TMX-induced KO of dynamins was efficient) from cells that did not respond to 4OH-TMC and internalized levels of Tf-A647 comparable to those of vehicle Ctrl treated cells (indicated as WT in the image). Scale bar= 20 μm, nuclei stained with Hoechst (red). C) Quantification of Tf-A647 uptake shown in A using automated image analysis. A threshold of Tf-A647 fluorescence intensity in the perinuclear area was set to distinguish WT from KO cells. Values represent the mean of three replicas. Error bars represent the STDEV. Statistical analysis was performed using an unpaired double tailed t-test (n.s.= non-significant).

**Figure S2.**
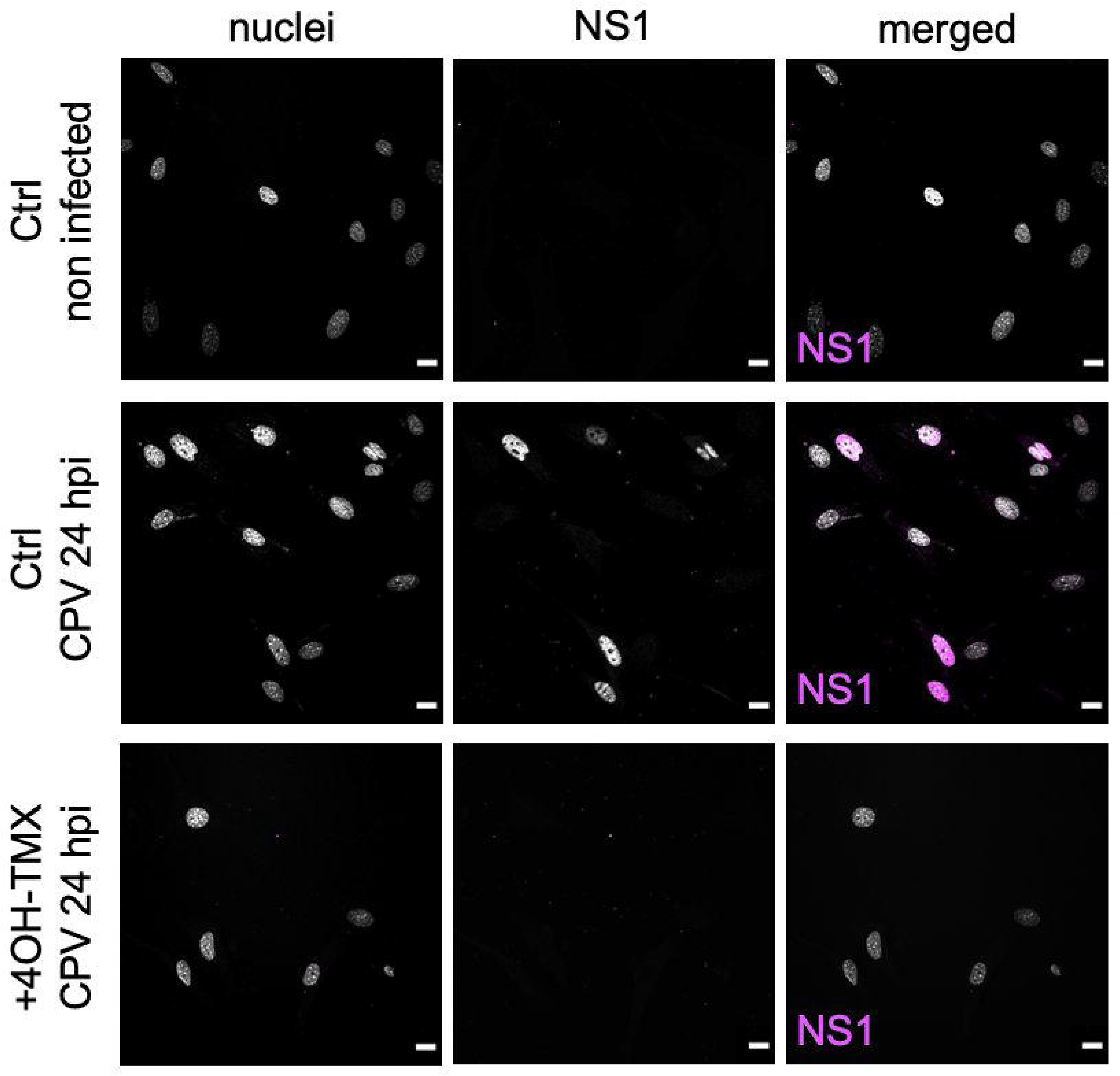
Dynamin 1,2 depletion blocks infection of CPV in MEF^DNM1,2 DKO^ cells. Representative confocal fluorescence images of MEF^DNM1,2 DKO^ cells infected with CPV for 24 h after 5 days treatment with EtOH vehicle control (Ctrl) or 4OH-TMX to induce dynamin depletion. Infected cells are visualized by immunofluorescence using an antibody against the viral non-structural protein NS1. Nuclei are visualized with DAPI DNA stain. Scale bar= 10 μm.

**Figure S3.**
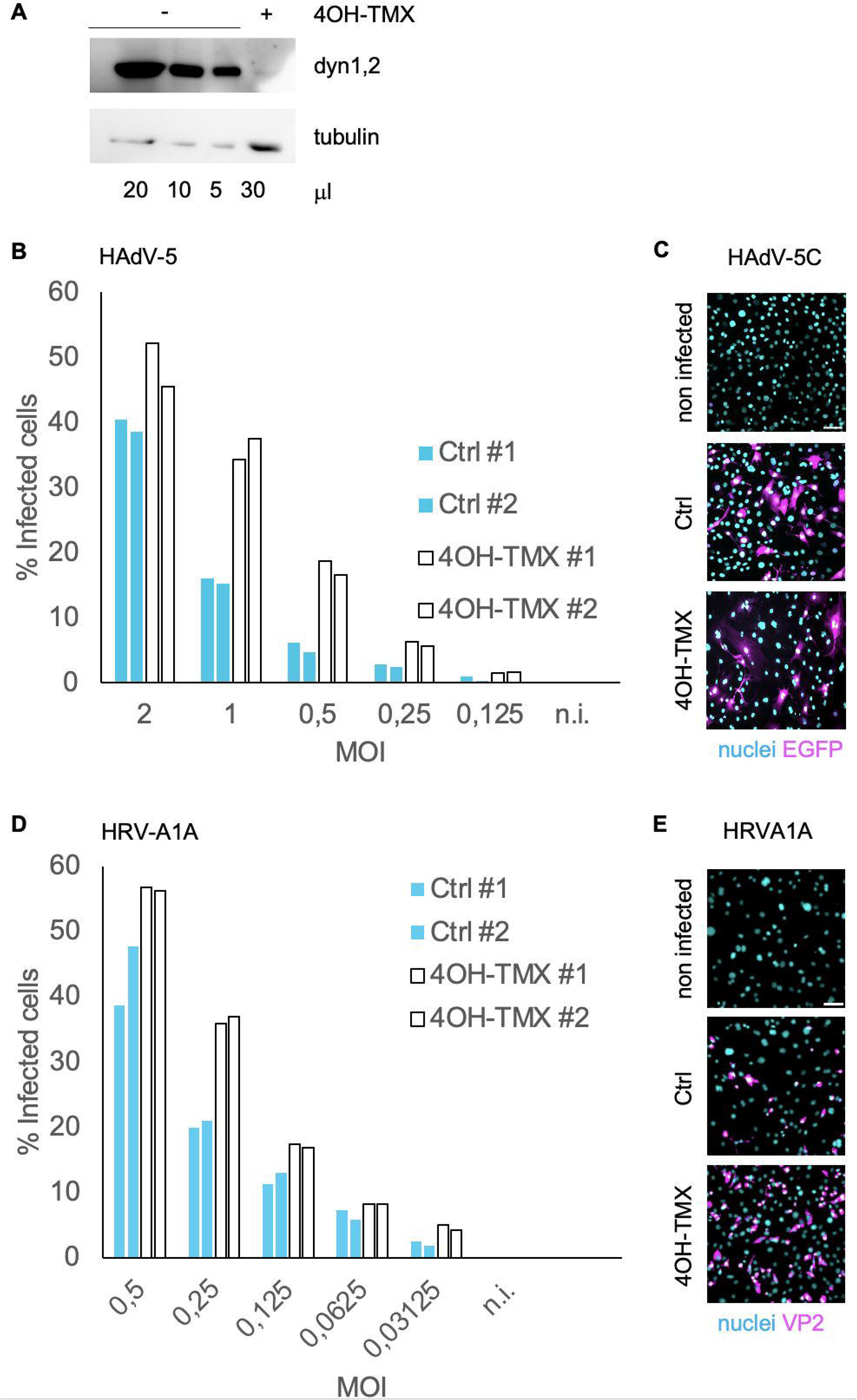
Dynamin depletion enhances infection of HAdV-5C and HRV-A1 in MEF^DNM1,2 DKO^ cells. A) Western blot analysis of dynamin 1,2 levels in MEF ^DNM1,2, DKO^ cells treated with vehicle control or 40H-TMX for 6 days. Indicated amounts of samples were loaded in the gel before blotting. Tubulin was used as a loading control. The Dyn1,2 antibody used recognizes both dynamin 1 and 2. B-C) Quantification of infection in MEF DNM1,2, DKO cells treated with vehicle control (Ctrl) or 4OH-TMX for 6 days and infected with AdV-5C for 22 h, and D-E) HRV-A1 for 24 h. Virus infection was determined by direct fluorescence imaging of HAdV-5C induced EGFP (C, magenta), and after immunofluorescence staining of HRV-A1 VP2 protein (E, magenta). Representative epifluorescence images are shown in C and E. The experiment was performed with two technical replicates (#1 and #2). n.i.= non infected. Scale bar= 10μm

**Figure S4.**
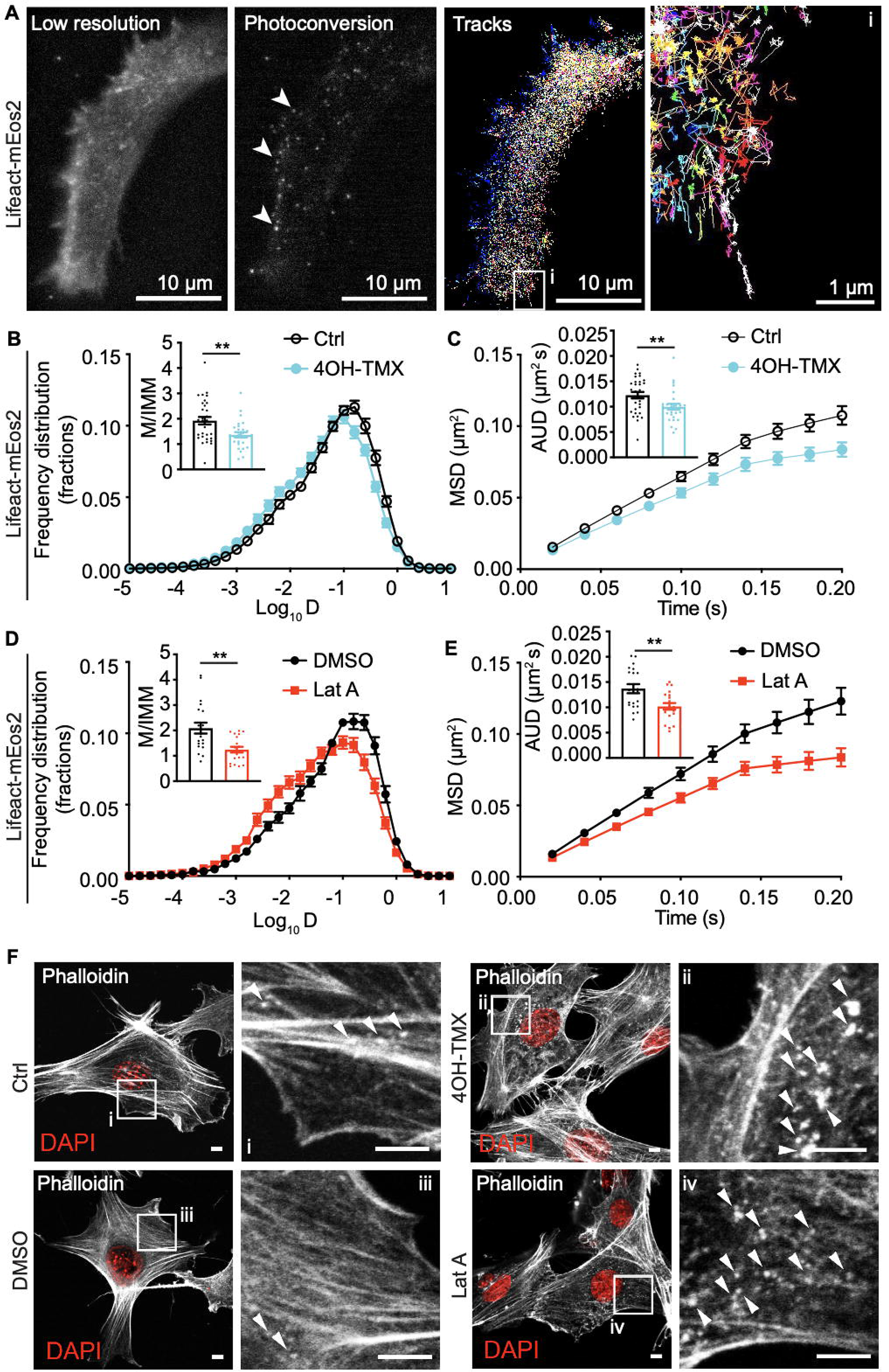
Dynamin-depletion induces slower actin dynamics. A) Super-resolution imaging using LifeAct-mEOS2. Oblique illumination with a low dose of UV light stochastically photo-converts the mEos2 protein from green to red fluorescence emission, allowing single particle identification and motion tracking in live cells. Arrowheads indicate diffraction limited fluorescence spots corresponding to single LifeAct-mEos2 molecules. The particle tracks identified during live cell imaging are indicated in the right panels. Each track line corresponds to the motion of a single LifeAct-mEos2 molecule. The white boxed area in Tracks image is expanded in i. Scale bar 10 μm. B-C) Quantification of single-molecule mobility of Lifeact-mEos2 in MEF^DNM1,2 DKO^ cells treated with vehicle control or 4OH-TMX for 6 days at 37°C. D-E) Quantification of single-molecule mobility of lifeact-mEos2 in MEF^DNM1,2 DKO^ cells treated with low dose Lat-A (0.1 µM) for 15 min at 37°C. MSD = mean square displacement. Values represent the mean of 28 Ctrl cells and 25 4OH-TMX treated cells in B, C and 19 Ctrl cells and 20 LatA treated cells in D, E. The error bar represents the standard error of the mean (SEM). Statistical significance was calculated by a Mann-Whitney test (**P<0.01). F) Immunofluorescence analysis of MEF^DNM1,2 DKO^ cells treated with EtOH vehicle control or 4OH-TMX for 6 days at 37°C, or with low dose Lat-A (0.01 mg/ml) for 10 min at 37°C. After fixation, actin fibers were stained with fluorescently conjugated phalloidin (white) and nucei were stained with the DNA dye DAPI (red). Arrowheads indicate punctated actin spots. Scalebar 1 μm.

**Figure S5.**
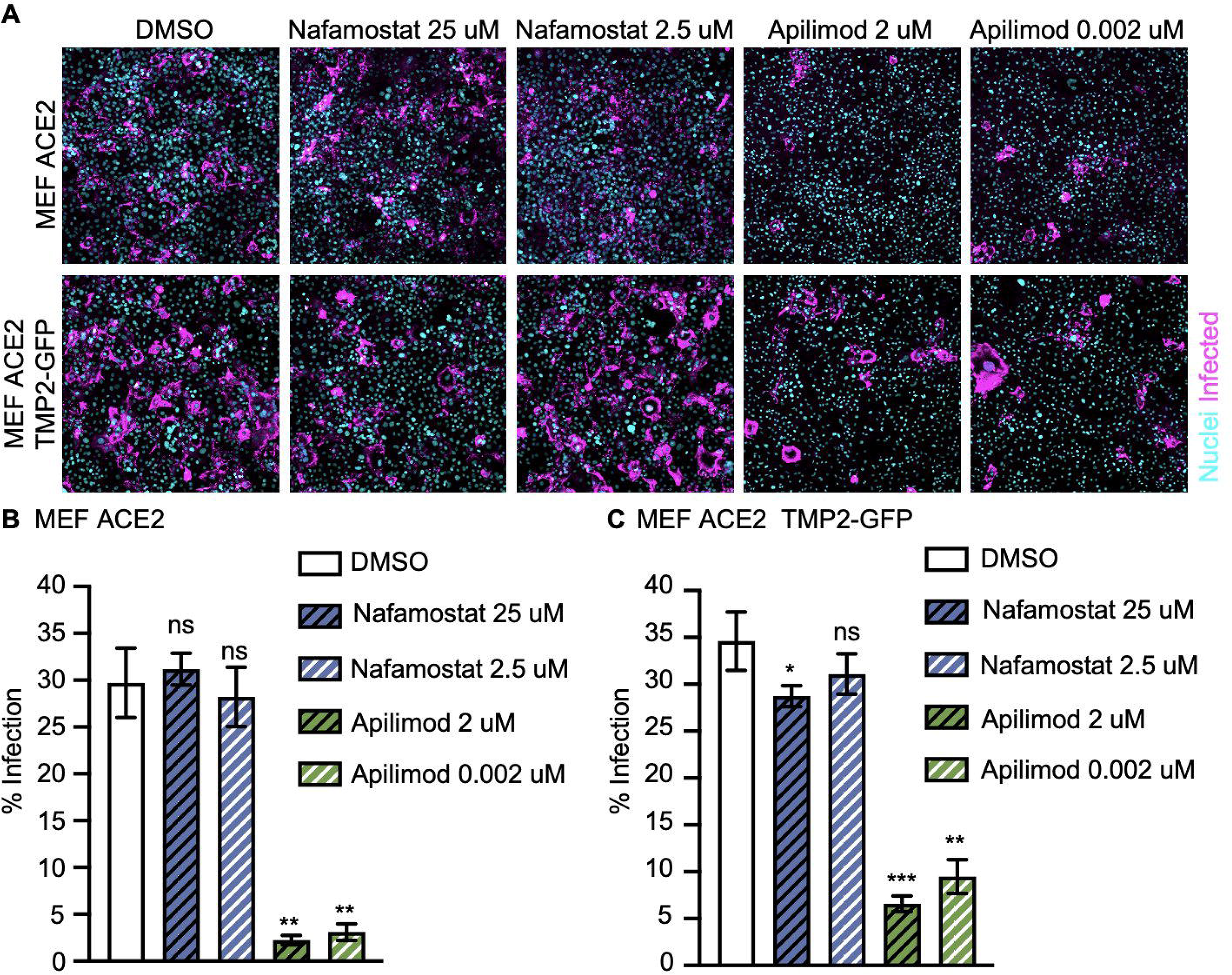
Sensitivity of SARS-CoV-2 ifection to inhibitors of endosome maturation and serine proteases. A) Representative fluorescence images of MEF^DNM1,2,3 TKO^ ACE2 and MEF^DNM1,2,3 TKO^ ACE2-TMPRSS2-GFP pretreated with nafamostat (25 μM or 2.5 μM), apilimod (2 μM or 0.002 μM) and infected with SARS-CoV-2 Wuh. B-C) Quantification by image analysis of the experiment shown in A. Values indicate the mean of at least three independent experiments and the error bars represent STDEV. Statistical significance was calculated using unpaired two-tailed t-test (*p<0,05; ** p<0,01; ***p<0,001; n.s.= non-significant).

